# A mechanically regulated liquid-liquid phase separation of the transcriptional regulator Tono instructs muscle development

**DOI:** 10.1101/2023.08.27.555003

**Authors:** Xu Zhang, Jerome Avellaneda, Maria L. Spletter, Sandra Lemke, Pierre Mangeol, Bianca H. Habermann, Frank Schnorrer

**Author notes:** equal contribution (Xu Zhang, Jerome Avellaneda).

## Abstract

Muscle morphogenesis is a multi-step program, starting with myoblast fusion, followed by myotube-tendon attachment and sarcomere assembly, with subsequent sarcomere maturation, mitochondrial amplification and specialisation. The correct chronological order of these steps requires precise control of the transcriptional regulators and their effectors. How this regulation is achieved during muscle development is not well understood. In a genome-wide RNAi screen in *Drosophila*, we identified the BTB-zinc finger protein Tono (*CG32121*) as a muscle-specific transcriptional regulator. *tono* mutant flight muscles display severe deficits in mitochondria and sarcomere maturation, resulting in uncontrolled contractile forces causing muscle atrophy during development. Tono protein is expressed during sarcomere maturation and localises in distinct condensates in flight muscle nuclei. Interestingly, internal pressure exerted by the maturing sarcomeres deforms the muscle nuclei into elongated shapes and changes the Tono condensates, suggesting that Tono senses the mechanical status of the muscle cells. Indeed, external mechanical pressure on the muscles triggers rapid liquid-liquid phase separation of Tono utilising its BTB domain. Thus, we propose that Tono senses high mechanical pressure to in turn adapt muscle transcription specifically at the sarcomere maturation stage. Consistently, *tono* mutant muscles display specific defects in a transcriptional switch that represses early muscle differentiation genes and boosts late ones. We hypothesise that a similar mechano-responsive regulation mechanism may control the activity of related BTB-zinc finger proteins that, if mutated, can result in uncontrolled force production in human muscle.

## Introduction

Muscle cells, also called muscle fibers, house thousands of contractile mini-machines called sarcomeres, which produce the mechanical forces powering animal movements. Each sarcomere is a stereotyped unit with a highly ordered architecture, in which centrally located bipolar myosin filaments face their motor domains towards highly ordered actin filaments. These actin filaments are cross-linked at the Z-discs bordering each sarcomere. Actin and myosin filaments are stably connected by spring-like titin filaments, thus achieving a high molecular order ^1–4^. Specific muscle fiber types differ in their transcriptional program resulting in specific differences in their sarcomeric composition and their biomechanical properties ^5–7^. As sarcomere architecture and its molecular components are conserved from flies to humans, *Drosophila* is a valid model to study sarcomere morphogenesis ^8,9^.

Striated muscle fibers form by a defined series of developmental events: myoblasts fuse to myotubes ^10^, which elongate and attach with both ends to tendon cells ^11,12^. After stable attachment, myotubes convert to myofibers by assembling immature myofibrils ^9,13–16^. These immature myofibrils consist of chains of short, immature sarcomeres, which are about 1.2 µm long in young zebrafish muscles ^17,18^ and 1.8 µm long in early *Drosophila* flight muscles ^15,19^. Next, these sarcomeres mature and grow to a length of about 3.4 µm in adult flight muscles ^19,20^. During this maturation phase, each sarcomere incorporates large amounts of specific sarcomere proteins ^19^, which adapt a high molecular order ^3^, resulting in the remarkable regularity of mature sarcomeres. Defects in sarcomere maturation can cause uncontrolled muscle contractions and severe muscle atrophy in vertebrates ^21–23^ and flies ^19,24–27^. Hence, correct sarcomere maturation is critical for regulated muscle force production.

In parallel to sarcomere morphogenesis, muscle fibers reorganise or generate various other internal structures essential for muscle function. Fibers assemble the postsynaptic part of the neuromuscular junction ^28^, position their nuclei equally throughout the fiber ^29,30^, and importantly gain a large number of mature mitochondria^31–34^, which match the specific metabolic needs of each particular muscle fiber type ^35,36^. All these internal structures are densely packed, resulting in high internal pressure in the developing muscles, with the myofibrils pushing the growing mitochondria into elongated shapes ^34^. How developing muscles coordinate and accomplish all these processes correctly in space and time is not well understood.

Here, we use the *Drosophila* indirect flight muscles to gain insight into how the transcriptional program of developing muscle fibers is dynamically adjusted to match the precise needs of each particular developmental stage. We have previously shown that sarcomeres develop in a very ordered sequence in flight muscles. Following assembly of immature sarcomeres into myofibrils at 30 h after puparium formation (APF) ^37^, additional immature sarcomeres are added to the existing myofibrils until about 48 h APF, but each sarcomere remains short and rather thin. After 48 h APF, all sarcomeres mature and grow in length and thickness to reach the adult length of about 3.4 µm ^3,19,20^. The sarcomere maturation phase is initiated by a transcriptional switch that down-regulates a large number of genes after 30 h APF, whereas other genes, including many sarcomere and mitochondria protein-coding genes, are strongly up-regulated ^19^. This switch needs to be precise, as even reducing levels of single sarcomere proteins can result in dominant flightless phenotypes ^38–40^ or flight muscle atrophy ^27^. Heterozygous loss of function mutations in individual sarcomere proteins can also cause dominant cardiomyopathies in humans ^41,42^. Thus, the expression levels of the various sarcomere components must be tightly controlled during sarcomere morphogenesis.

Here, we identified a specific role for *tono* (*CG32121*), a member of the conserved BTB-zinc finger family of transcriptional regulators, during indirect flight muscle morphogenesis. *tono* mutant flight muscles show a defective transcriptional switch after sarcomere assembly, resulting in severe sarcomere and mitochondria defects. Tono protein is present in muscle nuclei in small, mechanosensitive condensates which fuse in response to acute mechanical pressure. Thus, we propose that Tono senses an increase in mechanical pressure during mid-development to adapt the transcriptional status of the muscle to the developmental stage. This transcriptional adaptation contributes to correct sarcomere and mitochondria maturation.

## Results

### The BTB-Zinc finger protein *tono* is required for flight muscle morphogenesis

A genome-wide muscle-specific RNAi screen identified a potential role for *CG32121* for normal flight muscle development or maintenance ^43^. This gene was also found to be expressed in a muscle-specific manner in the recent adult fly cell atlas ^44^. *CG32121* is located on the third chromosome and encodes a BTB-zinc finger protein, for which no classical mutants were available (Figure 1A). We used our established TALEN and CRISPR protocols ^45,46^ to generate two *CG32121* deletion alleles (Figure 1A). We gave *CG32121* the name *tono,* for the phenotypic characteristics described below. The TALEN-induced *tono[1]* allele contains a 363 bp deletion removing the splice acceptor of the second exon and a large part of the BTB domain. In the CRISPR-induced *tono[2]* allele, we exchanged most of the *tono* locus with a STOP cassette and a 3xP3-dsRed marker leaving only 38 amino acids of the original Tono protein, which is likely a null allele (Figure 1A). Both *tono* alleles are homozygous viable but produce flightless animals, either homozygous or trans-heterozygous over a large deficiency, showing that *tono* is essential for flight muscle function (Figure 1B).

**Figure 1:**
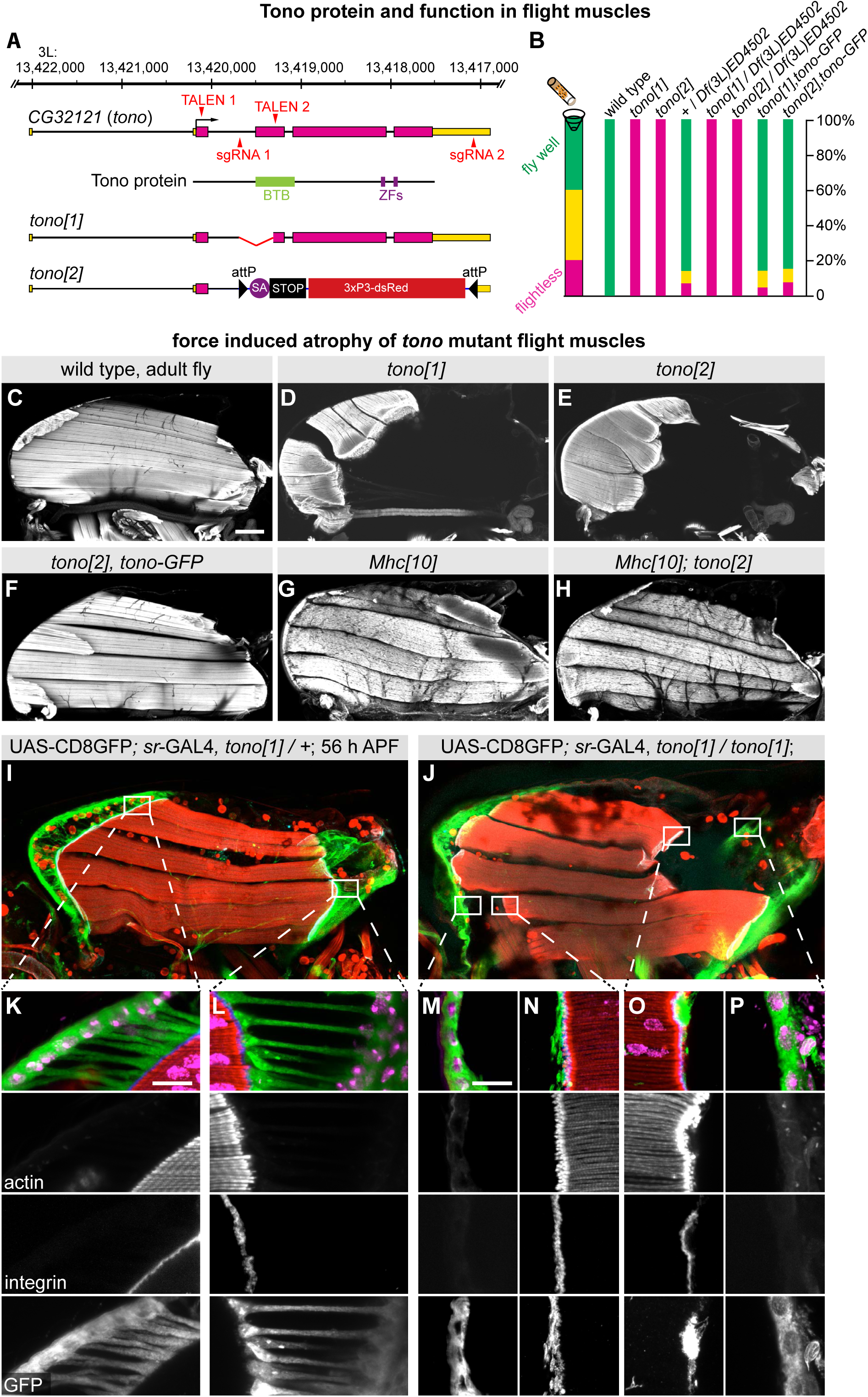
*tono* gene structure and adult flight muscle phenotypes. **(A)** Tono gene and protein structure, including the TALEN and sgRNA target sites used to generate the TALEN-induced *tono[1]* and the CRISPR-induced *tono[2]*, as well as the molecular nature of the alleles. (**B**): Flight test of the indicated genotypes. Note the complete flightlessness of both *tono* alleles. (**C-H**): Adult hemi-thoraces of wild type (C), *tono[1]* (D), *tono[2]* (E), *tono[2], tono-GFP* tagged Fosmid (F), *Mhc[10]* (G) and *Mhc[10]; tono[2]* (H) stained with rhodamine phalloidin to label F-actin. Note the severe flight muscle atrophy in *tono* alleles that is rescued in *Mhc[10]; tono[2]*. (**I-P**) Overview of heterozygous (I) and homozygous *tono[1]* pupa at 56 h APF expressing CD8-GFP in all tendon cells (J) stained with phalloidin in red, β-integrin in blue, GFP in green and nuclei in magenta. High magnification of the same genotypes at the anterior (K) or posterior tendon (L) of heterozygous or homozygous *tono[1]* (M-P). Note that all β-integrin remains at the ruptured muscle fiber ends together with tendon cell fragments in *tono[1]*. Scale bars represent 100 µm in C-J and 10 µm in K-P.

Histological analysis of the indirect flight muscles in *tono* homozygous or trans-heterozygous mutants showed that *tono* is required for flight muscle formation or maintenance, as young adult mutants displayed a severe flight muscle atrophy (Figure 1C-E, Supplemental Figure 1A-D). Importantly, this muscle atrophy phenotype could be rescued by re-expressing Tono-GFP under endogenous control using a GFP-tagged genomic fosmid construct ^47^ (Figure 1F, Supplemental Figure 1E, F), demonstrating that the *tono* loss of function phenotype is specific and that the *tono-GFP* fosmid is functional. The *tono* phenotype can also be rescued by re-expressing *tono-HA* in the developing flight muscles from around 24 h APF (Supplemental Figure 1H-J). Together, these data demonstrate that the BTB-Zn-finger *tono* is required specifically in muscles to prevent flight muscle atrophy.

### *tono* mutant muscles produce abnormally high forces during development

To identify the origin of the muscle atrophy in *tono* mutants, we investigated indirect flight muscle development during pupal stages. We found that *tono* mutant flight muscle fibers look normal until 48 h APF and are normally innervated by their motor neurons (Supplemental Figure 2A, B, F, J, N-P). However, from 56 h APF onwards, *tono[1]* flight muscles progressively detach. Detachment starts primarily at the posterior attachment sites (Supplemental Figure 2B-I), whereas *tono[2]* flight muscles are observed to stretch and display severe muscle thinning, eventually resulting in muscle rupture (Supplemental Figure 2J-M).

To investigate the detachment phenotype more closely, we labelled the tendon cell membrane with CD8-GFP. At 56 h APF, the wild-type tendon cells show the described, force-induced straight extensions from their basal side to the ends of the muscle fibers, where the integrin containing attachment sites are located (Figure 1I, K, L) ^37^. Surprisingly, in the detached *tono[1]* mutant muscle fibers, tendon cell membranes remained at the detached muscle ends (Figure 1J, M-P). This demonstrates that the tendon extensions themselves (and not the muscle-tendon junction) have been ruptured and torn from the tendon cell body. As this phenotype can be rescued by muscle-specific expression of *tono* (see Supplemental Figure 1), these data provide strong evidence that the forces produced in *tono[1]* flight muscles after 48 h APF are too high to be counteracted by normal tendon cells, resulting in tendon cell rupturing and muscle atrophy. To test this muscle force-induced tendon rupture hypothesis directly, we crossed the *tono* mutants into an *Mhc[10]* background, which lacks the force-producing Mhc isoform normally present in flight muscles ^3,48^. Indeed, the muscle atrophy phenotype is entirely rescued in *tono, Mhc[10]* double mutants (Figure 1G, H, Supplemental Figure 1G), demonstrating that fiber tearing and detachment are caused by uncontrolled high sarcomere forces in *tono* mutants.

### Tono is required for sarcomere and mitochondria maturation

To better understand the cause of the ectopic force production, we investigated sarcomere and mitochondria morphologies in developing *tono* mutant flight muscles. Until 48h APF, sarcomere morphology and length are comparable to wild type (Figure 2A-C, J; Supplementary Table 1). This shows that sarcomere assembly and addition, which are finished 48 h APF ^19^, do not require *tono* function. After 48h APF, wild-type sarcomeres mature and grow from 2 µm to about 2.9 µm in length at 72 h APF. However, *tono[1]* and *tono[2]* mutant sarcomeres fail to grow and remain about 2 µm long (Figure 2A-C, J, Supplemental Table 1). This growth defect is accompanied by actin accumulations in *tono[2]* myofibrils from 56 h APF onwards (Figure 2C). To investigate the functional consequences of this sarcomere maturation defect, we assayed the spontaneous twitching of flight muscles in living intact pupae. As reported before, we found that wild-type flight muscles display spontaneous twitching at 48 h APF, which ceases by 60 h APF when sarcomeres have adapted stretch-activation properties ^19^. However, we found robust twitches in *tono* mutant flight muscles at 60 h APF (Figure 2D, E; Supplemental Movies 1, 2). Together, these data demonstrate a severe sarcomere maturation defect in *tono* mutants that causes ectopic active forces in flight muscles, identifying abnormal myosin contractility as the likely cause of the muscle atrophy phenotype.

**Figure 2:**
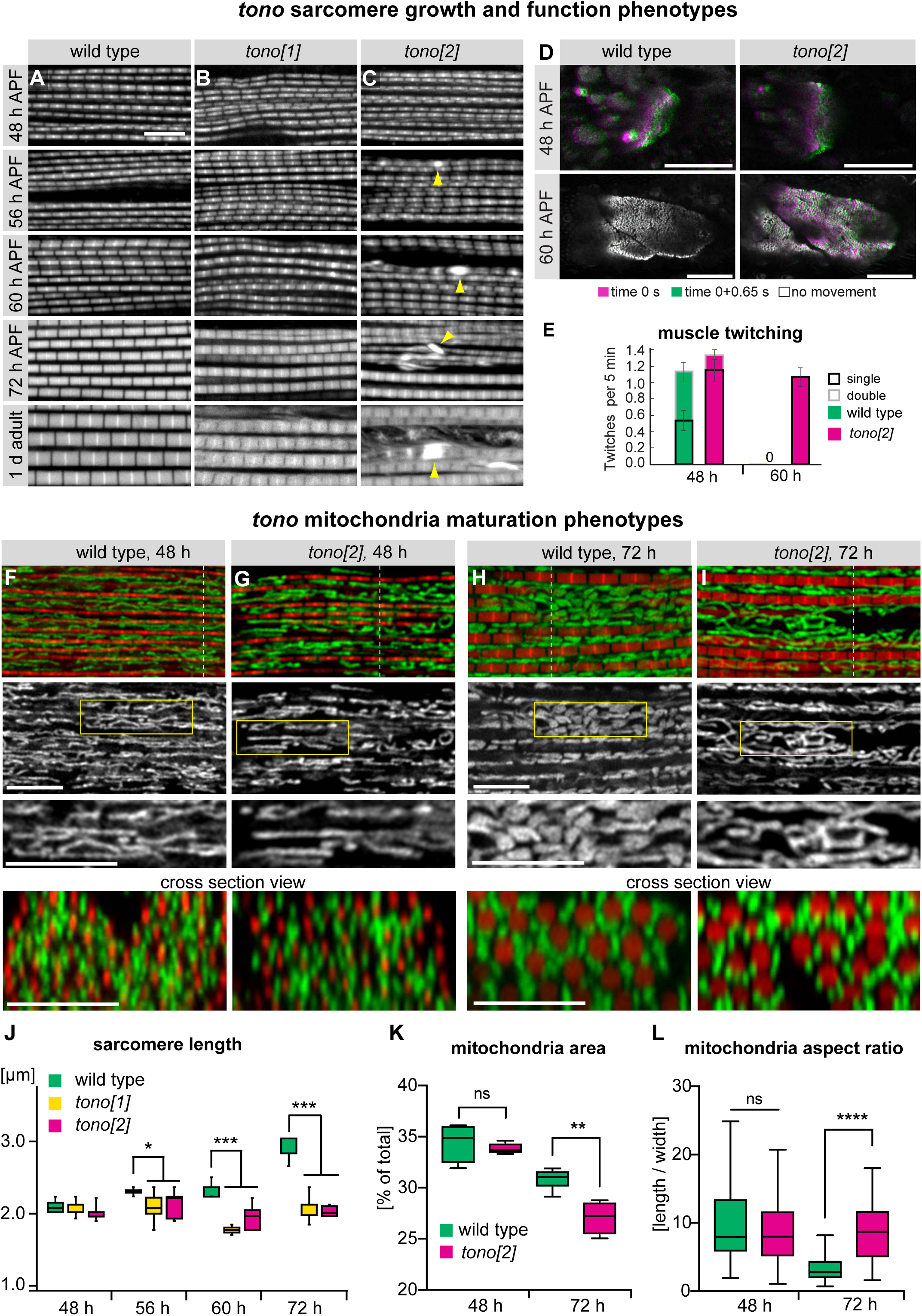
*tono* sarcomere and mitochondria phenotypes. (**A-C**) Developing myofibrils are visualised with phalloidin from wild-type (A), *tono[1]* (B) and *tono[2]* (C) mutant flight muscles at the indicated pupal time points. Note the large actin accumulations indicated by yellow arrowheads. Scale bar represents 5 µm. (**D, E**) Living wild-type or *tono[2]* mutant flight muscles at 48 h and 60 h APF (D). Two images taken at 0 seconds (magenta) and 0+0.65 seconds (green) were overlayed. White indicates no contraction; green and magenta colours show regions of contraction. Scale bars represent 50 µm. The frequencies of the muscle twitches were quantified per fiber per 5 min, error bars represent SEM (E). (**F-I**) Wild-type (F, H) and *tono[2]* mutant flight muscles (G, I) at 48 h and 72 h APF were stained for actin in read to visualise the myofibrils (phalloidin) and mitochondria in green (mit-GFP) and images with the LSM airyscan confocal. Yellow boxes show zoom-ins below. Cross-section views at the indicated dotted lines are shown at the bottom. Note that the 72 h *tono[2]* mitochondria look similar to mitochondria from 48 h APF wild-type muscles. Scale bars represent 5 µm. (**J-L**) Quantification of sarcomere length (J), mitochondria area (K) and mitochondria aspect ratio (L) from the above genotypes. ***P < 0.001, two-tailed unpaired Student’s t-test. Box-and-whisker plots show median as lines, 1^st^ quartile and 2^nd^ quartile as box and the min-max values in the whiskers.

In concert with the sarcomeres, the mitochondria network undergoes growth and rearrangement to fuel the intense contractions observed in mature muscle ^34^. By assaying mitochondria morphology, area and aspect ratio, we found that mitochondria in *tono* mutant flight muscles intercalated normally between the assembled myofibrils and show their normal elongated tube-like morphology at 48 h APF (Figure 2F, G, K, L). However, while wild-type mitochondria mature to shorter and fatter shapes at 72 h APF, *tono* mutant mitochondria maintain their thinner, elongated shapes that are typical of the earlier 48 h APF timepoint (Figure 2F-I). *tono* mutant mitochondria fail to change their aspect ratio and cover less area than in wild-type muscle, demonstrating their severe maturation defects (Figure 2K, L). Together, this demonstrates that *tono* is important for proper flight muscle maturation after 48 h APF.

### Tono is expressed in body muscle nuclei

The dramatic defects observed in *tono* mutant flight muscles made us wonder whether Tono protein is indeed a transcriptional regulator, as suggested from its protein domain structure. To address this, we first analysed Tono expression using the functional Tono-GFP fosmid. We found that Tono-GFP is localised within the nuclei of adult body muscles, including indirect flight, jump and leg muscles. However, Tono-GFP expression is not detectable in visceral muscles or other cell types of the adult thorax (Figure 3A-D), consistent with the fly cell atlas mRNA expression data ^44^. This demonstrates that Tono is expressed specifically in striated body muscles.

**Figure 3:**
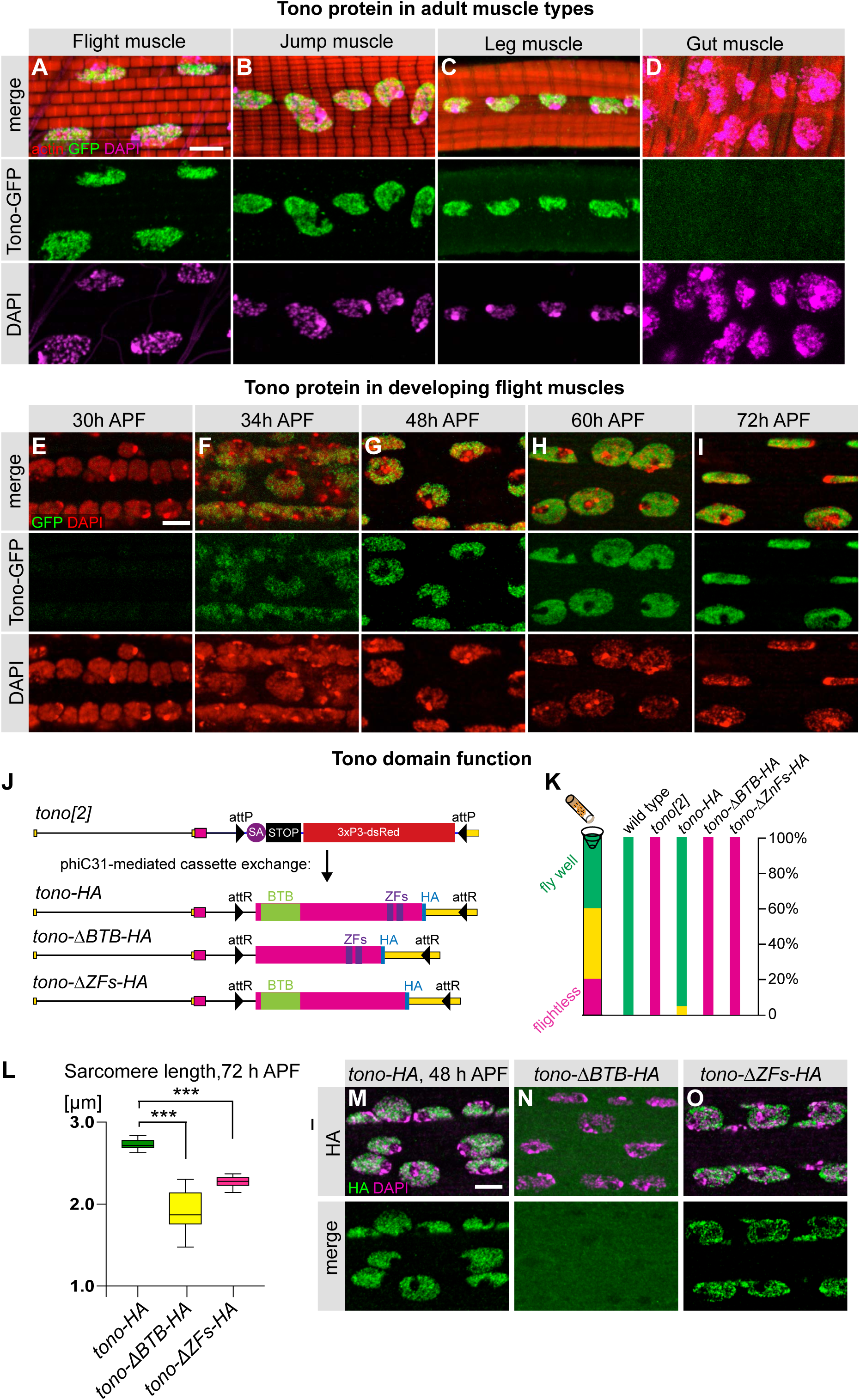
Tono protein expression and localisation. (**A-D**) Expression of Tono-GFP (fosmid) in different adult muscle types, dorsal-longitudinal flight muscle (A), jump muscle (B), leg muscle (C) and visceral muscle (D). Note that Tono-GFP localises to nuclei of all but the gut muscles. (**E-I**) Expression of Tono-GFP (fosmid) in developing flight muscles of the indicated stages. Note the high expression of Tono in nuclei after 34 h APF and its punctate localisation at 48 h APF. (**J-O**) Tono structure-function analysis. Scheme of the RMCE strategy creating the *tono-HA, tono-ΔBTB-HA* and *tono-ΔZfs-HA* alleles (J). Adult flight test (K) and sarcomere length in flight muscles at 72 h APF (L). ***P < 0.001, two-tailed unpaired Student’s t-test. Error bars represent the ± s.d. Localisation of Tono-HA (M), Tono-ΔBTB-HA (N) and Tono-ΔZfs-HA (O) in 48 h AFP flight muscles. Note the defect in nuclear localisation in Tono-ΔBTB-HA. All scale bars represent 5 µm.

As Tono has a role during muscle development, we investigated its mRNA and protein expression dynamics during pupal stages. We found that *tono* mRNA expression levels are induced from 16 h to 24 h and peak at 30 h APF in flight muscles (Supplemental Figure 3A) ^19^. Tono-GFP protein is barely detectable at 30 h APF when immature sarcomeres have just assembled (Figure 3E). At 34 h APF, Tono-GFP is present in indirect flight muscle nuclei and its expression strongly increases until 48 h, remaining high until 72 h APF (Figure 3F-I). Thus, Tono protein is expressed in the flight muscle just before the observed muscle maturation phenotypes are visible in *tono* mutants.

### Tono requires its BTB and Zn-finger domains

Having established the nuclear localisation of Tono, we wanted to investigate the roles of the BTB and Zn-finger domains. The CRISPR-generated *tono[2]* allele contains two flanking attP sites that can be utilised for phiC31 recombinase-mediated cassette exchange (RMCE) to engineer any designed *tono* allele ^46,49,50^. We inserted a *tono* cDNA tagged at the C-terminus with 2xHA, which will splice in frame with the remaining 38 amino acids of the endogenous first *tono* exon (*tono-HA*). We also inserted mutant *tono* cDNAs that are either lacking its BTB (*tono-ΔBTB-HA*) or zinc fingers (*tono-ΔZFs-HA)* (Figure 3J). All three HA-tagged proteins are expressed and the flies are homozygous viable. However, only the *tono-HA* flies can fly, whereas *tono-ΔBTB-HA* and *tono-ΔZFs-HA* are completely flightless (Figure 3K). We analysed the flight muscle morphology of these three different *tono* alleles and found that *tono-HA* flight muscles are normal and their sarcomeres grow normally (Figure 3L, Supplemental Figure 3B, E, H; Supplemental Table 1). In contrast, both *tono-ΔBTB-HA* and *tono-ΔZFs-HA* alleles display the typical myofiber rupture phenotype of *tono* null alleles at 72 h APF, with severe sarcomere defects (Figure 3L, Supplemental Figure 3B-J; Supplemental Table 1). This demonstrates that Tono requires both its BTB and zinc finger domains to support sarcomere maturation and flight muscle fiber integrity during muscle development.

The BTB domain of BTB zinc finger proteins often mediates protein-protein interactions, whereas zinc fingers are often implicated in binding to DNA ^51^. We investigated the distribution of Tono full-length and Tono deletion proteins during indirect flight muscle development and found that Tono-ΔBTB-HA is defective in nuclear enrichment. It appears to be able to enter the muscle nuclei but possibly is not retained there, resulting in an equal distribution between cytoplasm and nuclei at 48 h APF (Figure 3M, N). In contrast, Tono-ΔZFs-HA is normally localised in the muscle nuclei (Figure 5M, O), consistent with the interpretation that Tono’s zinc fingers are not required for nuclear localisation but rather for its function, possibly for its interaction with DNA.

### Tono regulates a metabolic switch during muscle maturation

To test if Tono regulates transcription during muscle development, we dissected indirect flight muscles from wild type and *tono[1]* mutants at 30 h and 48 h APF and performed transcriptomics analysis. In accordance with the wild-type morphology and the low expression of Tono at 30 h APF, we found few changes at 30 h APF, but a large number of genes (>2,000 genes) were differentially expressed at 48 h APF in *tono[1]* compared to wild type flight muscles (Supplemental Figure 4A, B Supplemental Data 1), This demonstrates that Tono is indeed a transcriptional regulator during muscle maturation stages.

We previously showed that developing wild-type flight muscles undergo a dramatic transcriptional switch between 30 h and 48 h APF, with more than 3,000 genes being up or down-regulated ^19^, which we reproduced here (Supplementary Data 1). Comparing wild type to *tono* mRNA-Seq data revealed that this transcriptional switch is impaired in *tono* mutant muscles, with many genes being less down-regulated or less up-regulated (Figure 4A). This difference in expression between wild-type and *tono* is particularly pronounced for genes that had been previously assigned to developmental clusters that change expression between 30 h and 48 h APF in wild type (Figure 4B, Supplemental Figure 5) ^19^. This is consistent with Tono being a transcriptional regulator with an important role at this key stage of muscle development.

**Figure 4:**
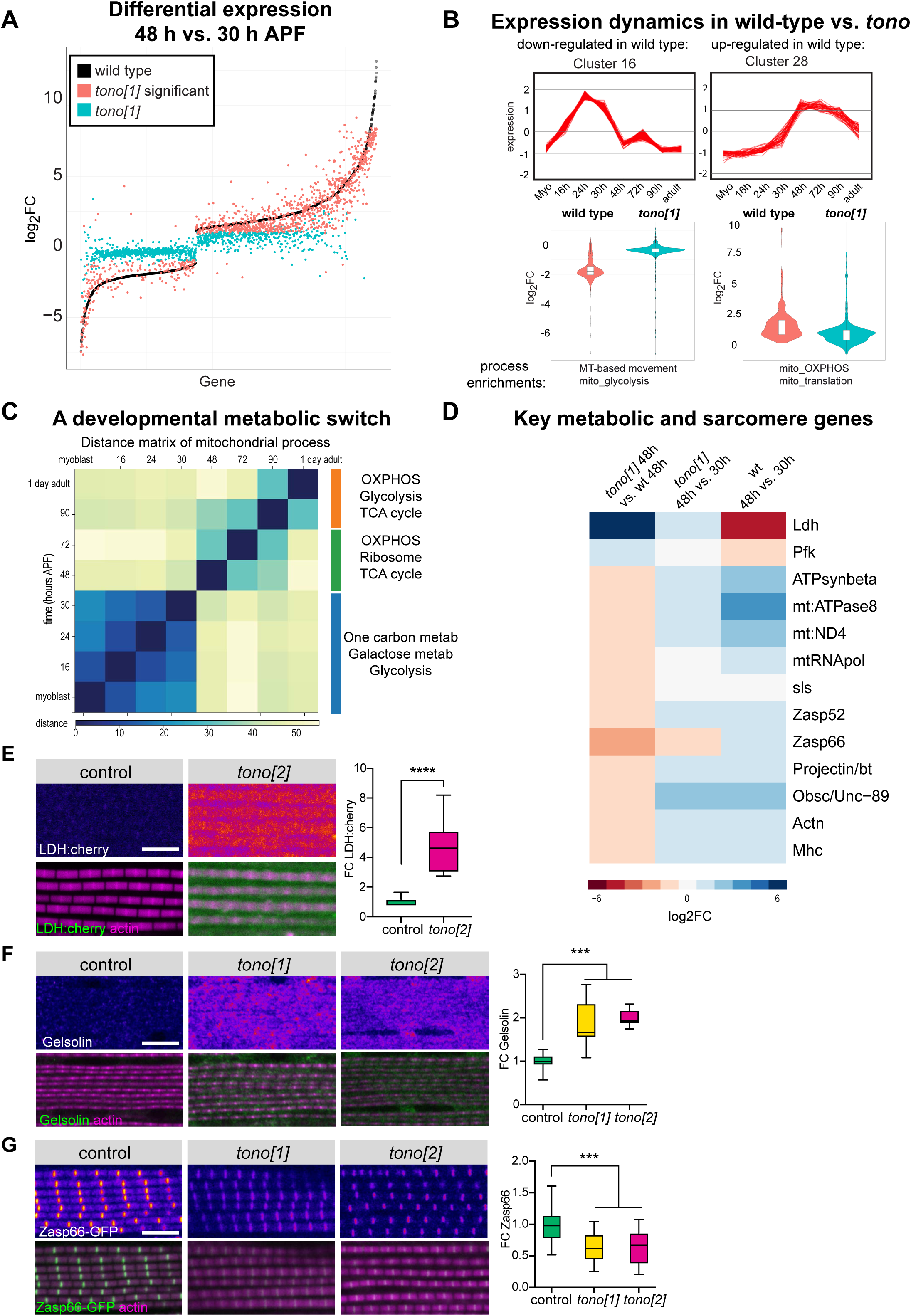
Tono regulates a transcriptional switch during muscle development. **(A)** Black dots represent 2,238 significantly changed genes comparing wild type 30 h to 48 h APF flight muscles. Note that the same comparison in *tono[1]* results in many genes that are above the left part of the curve (less down-regulated) or below the right part of the curve (less up-regulated). Significant differences between 30 h to 48 h in *tono[1]* flight muscle are shown in blue. All data are listed in Supplemental Data 1. (**B**) Expression dynamics during flight muscle development; one selected cluster down-regulated at 30 h APF (Cluster 16) and one up-regulated cluster (Cluster 28) are shown with the enriched processes from ^19^. Violin plots show that the same genes have weaker expression changes in *tono[1].* (**C**) Applying Phasik ^53^ to the developmental transcriptomics data ^19^ revealed three distinct phases during development that change from 30 h to 48 h APF, and at 72 h to 90 h APF, respectively. (**D**) Log2FC in wild type and *tono[1]* at 48 h vs 30 h APF of key enzymes of the glycolysis, lactate metabolism and oxidative phosphorylation from the transcriptomics data. (**E-G**) Verification of the expression changes in control vs *tono[2]* flight muscles using a lactate dehydrogenase (LDH) reporter (E), anti-Gelsolin antibody staining (F) or Zasp66-GFP intensity (G). ***P < 0.001, two-tailed unpaired Student’s t-test. Box-and-whisker plots show median as lines, 1^st^ quartile and 2^nd^ quartile as box and the min-max values in the whiskers. Scale bars represent 5 µm.

In accordance with our phenotypic data demonstrating a mitochondrial defect, the developmental clusters 16 and 28 had been shown to be enriched for metabolic pathways, including glycolysis and oxidative phosphorylation (Figure 4B) ^19^. We found similar pathways, including TCA cycle and glycolysis, as being amongst the top *tono*-regulated KEGG pathways at 48 h (48 h *tono[1]* vs. wild type) as visualised by the EnrichR analysis tool (Supplementary Data 1). To explore the dynamics of the metabolic pathways in more detail, we used the mitoXplorer tool ^52^ and the Phasik method ^53^ and applied them to developing flight muscles using our previously published wild-type time course mRNA-Seq data ^19^. These methods identified a metabolic switch between 30 h and 48 h APF in developing wild-type muscles, with glycolysis and one carbon metabolism being dominant before 40 h APF and oxidative phosphorylation, TCA cycle and protein synthesis in mitochondria being dominant after 40 h APF (Figure 4C, Supplemental Data 2). This switch is impaired in *tono* mutant muscles. We found that the mitochondrial translation machinery is less upregulated at 48 h APF in *tono* compared to wild-type. Consequently, the mitochondria-encoded OXPHOS components, including mt:ND4L and mt:ATPase8, are strongly downregulated, as well as large numbers of the nuclear-encoded OXPHOS components, including ATPsynthase β (Figure 4D, Supplemental Figure 4C). In contrast, phophofructokinase (Pfk), the rate limiting enzyme of glycolysis, as well as the pyruvate processing enzyme lactate dehydrogenase (LDH), fail to be downregulated in *tono* mutant muscle (Figure 4D, Supplemental Figure 4C). We further confirmed the LDH down-regulation *in vivo* with a transcriptional LDH reporter line in flight muscles (Figure 4E). Together, these data demonstrate that Tono transcriptionally instructs the metabolic switch during mid-stages of flight muscle development.

The second large class of genes that change expression during mid-stages of muscle development code for microtubule and actin dynamics, as well as for sarcomere proteins, the latter of which are strongly induced ^19^. Again, we find that this switch is impaired in *tono* mutant muscles with the key sarcomere components Mhc, Actn, Zasp52, Zasp66, Unc89/Obscurin and the titin homologs bt/Projectin and sls being improperly upregulated (Figure 4D, Supplemental Data 1). We verified the increased expression of the actin-severing protein Gelsolin, which is normally down-regulated from 30 h to 48 h APF, by antibody stainings in *tono[1]* and *tono[2]* mutants (Figure 4F). We also verified that expression of the sarcomere Z-disc component Zasp66, whose expression normally increases from 30 h to 48 h, is reduced in *tono[1]* and *tono[2]* mutants by quantifying GFP fluorescence (Figure 4G). These transcriptional changes may explain the sarcomere growth defect and the uncontrolled high forces in *tono* mutant muscles.

### Tono localises to nuclear droplets

How does Tono contribute to this transcriptional change at 48 h APF? By inspecting Tono-GFP or Tono-HA nuclear localisation in detail, we discovered that Tono localises non-homogenously throughout the muscle nuclei in a speckled pattern that is reminiscent of nuclear granules or phase-separated condensates (Figure 5A). This pattern reminded us of stress granules that were reported *in vitro* to respond to osmotic stress ^54^. To test if the Tono localisation pattern also responds to osmotic stress, we incubated 48 h APF pupae for 30 min with PBS or PBS plus 250 mM NaCl, to mimic osmotic stress. Interestingly, we found that the small Tono-GFP droplets observed in directly fixed or PBS-treated muscle nuclei condense into large droplets after incubation in high salt (Figure 5A-C), demonstrating that the Tono-positive droplets can dynamically rearrange. These Tono-GFP droplets are preferentially located at the nuclear periphery (Figure 5B’, C’) and also form at later developmental stages upon osmotic stress or in response to sucrose-induced osmotic stress (Supplement Figure 6A-E). They are not enriched for the heterochromatin marker HP1 (Supplement Figure 6F, G), which forms liquid droplets in early fly embryos or human cells ^55,56^. To test which domain of Tono is needed for droplet formation, we tested our HA-tagged Tono deletion mutants and found that the BTB domain but not the Zn-fingers is needed for droplet formation at 72 h APF (Figure 5D-I). These findings are in accordance with Tono being a dynamic transcriptional regulator, whose activity may be adjusted according to the internal status of the muscle cell.

**Figure 5:**
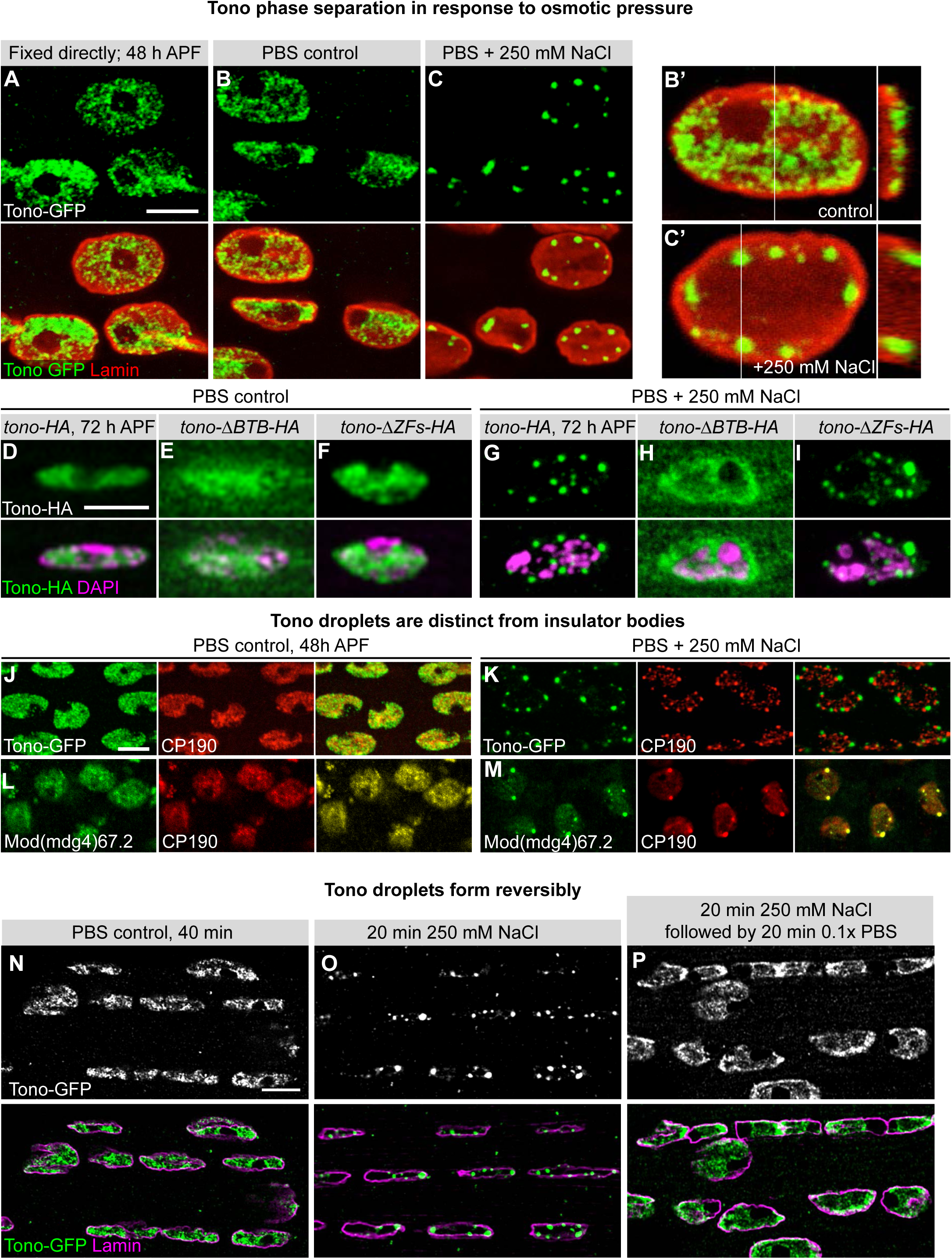
Tono forms liquid droplets in muscle cell nuclei. (**A-C**) Tono-GFP localisation pattern in flight muscles at 48 h APF either fixed directly (A) or incubated with isotonic (B, B’, same nucleus) or hypertonic (C, C’) solution for 30 min. GFP is shown in green and anti-Lamin in red. Note the large phase-separated Tono droplets after hypertonic treatment. (**D-I**) Tono-HA (D, G), Tono-ΔBTB-HA (E, H) and Tono-ΔZfs-HA (F, I) in 72 h AFP flight muscles after 30 min incubation in isotonic or hypertonic solution. Note that Tono-ΔBTB-HA does not form droplets. (**J-M**). Co-staining of Tono-GFP (green) with insulator complex components CP190 or Mod(mdg4)67.2 (red) in 48 h AFP flight muscles after 30 min incubation in isotonic (J, L) or hypertonic solution (K, M). Note that Tono droplets are distinct from insulator bodies. (**N-P**) TonoGFP droplets form reversibly. Compare TonoGFP 40 min in isotonic solution (N) or 20 min in hypertonic solution (O) to 20 min hypertonic solution, followed by 20 min hypertonic solution (P). All scale bars represent 5 µm.

### Tono droplets are distinct from insulator bodies

A similar dynamic re-localisation upon osmotic stress had been described *in vitro* for a class of proteins called insulator bodies, including CP190, BEAF-32, Su(Hw) and Mod(mdg4)67.2 ^54^. CP190 and Mod(mdg4)67.2 also contain BTB domains. As these components are expressed ubiquitously, we hypothesized that Tono may be a muscle-specific component of the insulator complex. However, we found that although all tested insulator components are recruited to large droplets upon osmotic stress in flight muscles, Tono-positive droplets are distinct from and do not overlap with droplets positive for CP190, BEAF-32, Su(Hw) or Mod(mdg4)67.2 (Figure 5J-M, Supplemental Figure 7A-G). These data show that Tono droplets are distinct from insulator bodies.

It is important to note that most of the generally accepted insulator complex components appear dispensable for normal *Drosophila* development. Null or strong hypomorphic alleles for *CTCF*, also a core insulator-complex member ^57^, *BEAF-32* or *CP190* are viable ^58–61^ or only pharate lethal and their flight muscles do not display gross defects (Supplemental Figure 7H-K). Thus, in contrast to the classical insulator complex components, Tono has an essential role in sarcomere and mitochondria maturation during indirect flight muscle development and may fulfil this role by altering its localisation pattern in response to the internal status of the muscle cell.

### Tono droplets form reversibly

The rapid formation of the large, round Tono droplets suggested a mechanism of droplet formation by liquid phase condensation. Such condensates form by a regulated liquid-liquid phase separation (LLPS) process, a widely accepted regulatory phenomenon in biology ^62–65^. LLPS droplets are characterized by their highly dynamic nature, their ability to fuse and the reversibility of droplet formation ^63^. Our experiments above demonstrated both the dynamic nature of Tono droplets as well as the ability of droplets to fuse to form larger condensates (Figure 5 A-C, Supplemental Figure 6 A-E). Another key feature of liquid condensates is that they form reversibly ^63^. We therefore tested if releasing the osmotic stress can result in disassembly of Tono-positive droplets. Indeed, we found that incubation in hypo-osmotic conditions results in droplet disassembly (Figure 5N-P). Tono-positive droplets therefore meet all three criteria – fusion, dynamic and reversible – to be classified as LLPS droplets.

### Tono droplets respond to forces

What could be the endogenous function of the Tono droplets? We hypothesised that our experimentally applied osmotic stress might model a naturally occurring phenomenon during flight muscle development. Just as osmotic stress transports water out of cells and thus intensifies the internal crowding and intracellular pressure, the radial growth of myofibrils and mitochondria generates crowding and thus increases internal pressure in the developing flight muscles, as we have shown in adult flight muscles ^34^. To test if such a high pressure might be present at 48 h APF and possibly sensed by Tono in the nuclei, we quantified nuclear shapes in flight muscles and found that flight muscles have strongly elongated, flat disc-like shaped nuclei that are squeezed between the bundles of growing mitochondria and myofibrils (Figure 6A). Compared to the less squeezed jump muscle nuclei, the flight muscle nuclei are more elongated (Figure 6A, B, D), suggesting a particularly high pressure on the flight muscle nuclei. This pressure is largely produced by the assembled myofibrils, as the nuclei are more round in *Mhc[10]* mutants, which display defective myofibrils (Figure 6C, D).

**Figure 6:**
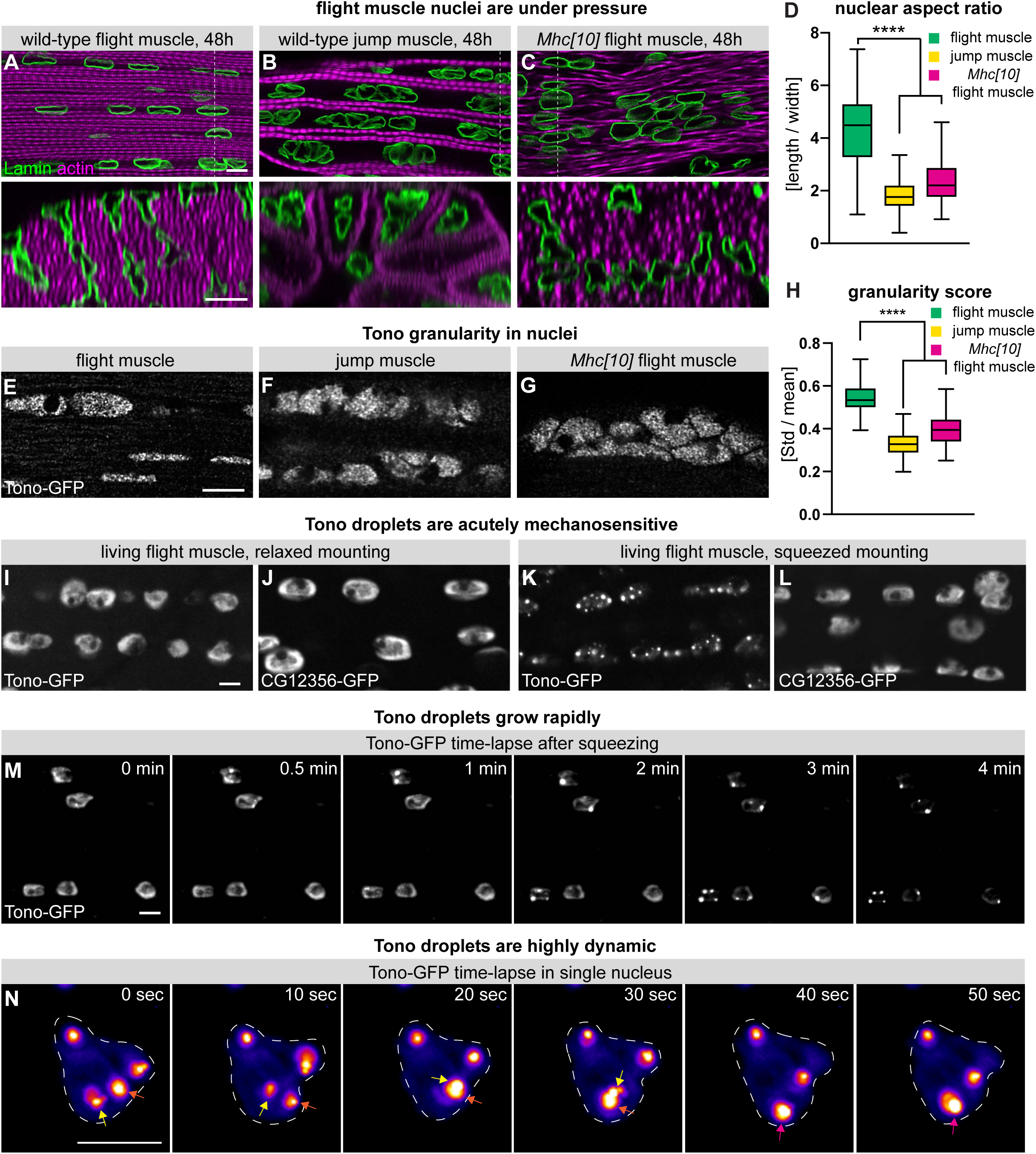
Tono droplets are acutely mechanosensitive. (**A-C**) 48h APF wild-type flight muscle (A), jump muscle (B) and *Mhc[10]* mutant flight muscle nuclei were stained with anti-Lamin (green) and phalloidin (magenta). Cross-section views at the dotted lines are shown below. Note the elongated nuclei in wild-type flight muscles. (**D**) Quantification of nuclear aspect ratios. (**E-H**) 48h APF wild-type flight muscle (A), jump muscle (B) and *Mhc[10]* mutant flight muscle nuclei expressing Tono-GFP were stained with GFP-nanobody and Tono granularity was quantified (H). (I-L) Live imaging of adult flight muscles expressing Tono-GFP and CG12356-GFP mounted relaxed (I, J) or squeezed (K, L). Live imaging of Tono-GFP granules after squeezing (see also Supplemental Movie 3). High-speed live imaging of one nucleus expressing Tono-GFP granules after squeezing (see also Supplemental Movie 4). Note that 2 neighbouring granules marked by yellow and orange arrows fuse (red arrow). All scale bars represent 5 µm.

To test if the small Tono-GFP droplets respond to pressure on the muscle nuclei, we quantified the Tono-GFP granularity by analysing the Tono-GFP intensity distributions in the different nuclei. The concentration of Tono in large droplets would result in a larger standard deviation of Tono intensities in the nuclei, compared with more homogenously distributed small droplets (Supplemental Figure 8, see methods for details). Using these criteria, we found that the granularity in wild-type 48 h APF flight muscles is larger than in jump muscles or *Mhc[10]* mutant flight muscles (Figure 6E-H). As the forces on the nuclei are lower in jump muscles or *Mhc[10]* mutant flight muscles compared with wild-type flight muscles, these results strongly suggest that Tono droplets are mechanosensitive and thus may sense the internal pressure of the developing flight muscles. Hence, we chose the name Tono, inspired by the tonometer used to measure the pressure within the human eye.

### Tono droplets respond to acute force changes

Osmotic pressure is reported to mimic cellular compression forces, however, it is often larger than hydrostatic pressure induced by the cytoskeleton ^66,67^. Hence, we wanted to directly increase the mechanical pressure and observe the consequences on Tono droplets. To do so, we mounted living Tono-GFP or CG12356-GFP expressing control flight muscles either “relaxed” with spacers, or “squeezed” with the coverslip slightly pressing on the flight muscles. Strikingly, we observed the formation of large Tono-GFP droplets after the squeezed mounting method, whereas control CG12356-GFP did not accumulate in droplets (Figure 6I-L), showing that Tono is selectively sensitive to the mechanical state of the muscle cell. As this mounting method is compatible with live imaging, we discovered that Tono-GFP is highly dynamic and phase-separates into large droplets within a few minutes after the pressure was applied (Figure 6M, Supplemental Movie 3). This demonstrates that Tono reacts to acute forces on the flight muscle nuclei by forming large droplets.

One important feature of liquid-phase separated droplets or condensates is their rapid fusion to larger round droplets ^63^. By recording high-speed high-resolution movies of the Tono droplets after mechanical squeezing, we indeed found that smaller Tono droplets fuse to large ones in less than 1 minute (Figure 6N, Supplemental Movie 4). Together, these results demonstrate that Tono can acutely sense the compression status of flight muscle nuclei and thus may adjust the transcriptional program of the muscle fiber to its developmental status.

## Discussion

Muscles are extremely protein-dense tissues that organise their internal organelles and contractile elements with pseudo-crystalline regularity ^68^. At the same time, their rigid myofibrils are under high mechanical tension along their long muscle axis ^15,16,69^. This results in a build-up of high internal pressure, with myofibrils squeezing mitochondria and nuclei into elongated shapes ^34,70^. However, early during muscle development, protein content and pressure are low, as myofibrils and mitochondria are both thin, and nuclei are round ^34^. Thus, it would be advantageous for a developing muscle fiber to sense when a certain level of pressure has been reached, as then the transcriptional program to promote myofibril and mitochondrial growth can be triggered ^19^.

Here, we identified the BTB Zn-finger protein Tono as an important regulator of this transcriptional switch during flight muscle development. In *tono* mutants, the normal developmental regulation of a group of genes coding for sarcomere and mitochondria components is defective. This results in too short sarcomeres that produce uncontrolled forces, causing muscle and tendon tearing and resulting in flight muscle atrophy. Related hypercontraction phenotypes are also caused by mutations in the splicing regulator Bruno or the myosin regulators Troponin I and Troponin T ^24,71–73^. However, in these cases muscle hypercontraction and ruptures only occur at the adult stage. Furthermore, *tono* mutant flight muscles display additional defects in their developmental metabolic switch, resulting in a failure of mitochondria growth. This demonstrates the uniqueness of the *tono* phenotype underscoring its key importance for normal muscle development.

How does Tono regulate the transcriptional switch? We propose that Tono senses the internal pressure state of the muscle cell in its nuclei to instruct transcription differentially during myofibril and mitochondria maturation. The supporting evidence for this hypothesis is fourfold. First, we found that the growing myofibrils and mitochondria squeeze the nuclei into elongated flat discs at this stage. Second, increasing the internal pressure acutely by hyperosmotic conditions results in a dramatic condensation of Tono protein into large, round droplets in the nuclei. This treatment induces water efflux from cells, and thus dramatically increases the jamming of all components ^74,75^, hence likely also the Tono concentration in the nuclei. Formation of these droplets is moreover reversible, demonstrating dynamic assembly and disassembly of Tono droplets. Third, acute mechanical pressure induces the fusion of small Tono droplets into larger droplets within a few minutes. This demonstrates the acute mechano-sensitivity of the Tono localisation pattern in the nuclei. As the range of mechanically induced hydrostatic pressure is believed to be 100 times lower than osmotically caused pressure differences (Mosaliganti et al., 2019; Torres-Sánchez et al., 2021; Venkova et al., 2022), this result underscores the high mechano-sensitivity of the Tono localisation pattern. Fourth and most importantly, the endogenous Tono localisation pattern is mechanosensitive as its granularity in the muscle nuclei decreases in response to a reduction of mechanical pressure in myosin mutants. Together, these data strongly suggest that Tono senses mechanical pressure in the flight muscle nuclei during development.

How does Tono droplet formation impact its activity? Tono requires its BTB domain to change its localisation in response to osmotic pressure, suggesting that it uses this domain to sense or respond to mechanical pressure. The dynamics of this change follows the typical fast fusion dynamics of smaller droplets to large ones, as described for various liquid-liquid phase separations ^63,65,76^. In particular in the nucleus, liquid-liquid phase separation has been proposed as a mechanism to subdivide the nucleus into different activity domains, with active and inactive DNA compartments, further subdivided into topologically associating domains (TADs) ^76,77^ and the formation of so-called super-enhancers that result in high expression rates ^78^. Established insulator proteins, like Cp190 and Mod(mdg4)67.2 also contain BTB Zn-finger domains ^77^ and condense into large droplets distinct from the Tono droplets under high osmotic pressure. What makes Tono special is that it is also mechano-sensitive and that it displays a muscle-specific loss of function phenotype. Hence, it is conceivable that Tono does impact chromosome organisation or DNA binding in response to mechanical pressure. However, currently it is unknown if and where the Tono Zn-fingers bind to DNA. As chromatin-immunoprecipitation experiments from flight muscles have proven difficult ^36^, solving this important question will require the development of additional tools in the future.

The nucleus is a prime organelle where multiple sensory inputs are integrated to monitor the mechanical status of a cell ^79^. It can directly sense cellular osmolarity in plant cells and hence responds to dehydration or mechanical impact by changing transcription ^80^. Deformation of the nuclear membrane in mammalian cells can also directly signal to the actomyosin cytoskeleton via the activation of a cytosolic phospholipase to for example allow rapid squeezing of migratory cells through fine pores ^81,82^. Particularly relevant for muscle cells is a MAP3K called ZAKβ that was recently shown to change its activity and localisation in the nucleus in response to compression or cyclic contraction in mammalian muscle cells ^66^. ZAKβ mutant mice display muscle nuclei position defects and show deficits in sarcomere protein expression and muscle maintenance ^66^. Furthermore, mutations in lamins or KASH-SUN proteins in the nuclear membrane that are implicated in mechano-sensing ^83,84^ cause severe Emery-Dreifuss muscular dystrophies ^85^. This underscores the importance of pressure sensing in muscle cells from their birth to their death.

Mammalian genomes contain about 50 BTB Zn-finger genes with no clear Tono homolog. However, mutations in the human BTB Zn-finger ZBTB42 result in lethal congenital contractures at birth. ZBTB42 is highly expressed in muscles, and zebrafish knock-down larvae show severe sarcomere and myofibril defects ^86^, suggesting that ZBTB42 might be a functional homolog of Tono in vertebrate muscles.

## Methods

### Fly strains and genetics

All fly work was performed at 27 °C under standard conditions unless specified. *tono[1]* and *tono[2]* were generated by TALEN and CRISPR-Cas9 mediated genome editing as described in the following section. The *CG32121* deficiency line (*w*^1118^*; Df(3L)ED4502, P{3’.RS5+3.3’}ED4502/TM6C, cu*^1^ *Sb^1^*) was obtained from the Bloomington *Drosophila* stock centre (BDSC). *tono-GFP* Fosmid (fTRG10059) was generated within the FlyFos collection ^47^. In the rescue experiments, *tono-GFP* Fosmid was recombined with *tono[1]* or *tono[2]* and the recombinant was identified by PCR. Tissue-specific rescue was performed at 18°C by driven UAS*-tono-HA* (obtained from Johannes Bischof ^87^) with the flight muscle-specific driver *Act88F-*GAL4 ^88^ in the *tono[1]* or *tono[2]* background. Hypercontraction rescue was performed with *Mhc[10]; tono[1] or Mhc[10]; tono[2]* flies by using the flight muscle-specific myosin splicing mutant *Mhc[10]* ^89^. Tendon cell labelling was performed by crossing UAS*-CD8GFP* with *stripe-*GAL4 ^90^ as control or UAS*-CD8GFP; tono[1]* with *tono[1]*, *stripe-GAL4*. Mitochondria were labelled by expressing a GFP fused to a mitochondria matrix peptide (mit-GFP) ^34^. The indirect flight muscles were labelled with *Mef2-*GAL4; UAS*-GFP-Gma* to facilitate the muscle dissection transcriptomics. Both *CTCF[GE24185]* and *CTCF[P30.6]* (from Rainer Renkawitz) ^57^ are strong alleles for *CTCF*. *BEAF-32[AB-KO]* ^91^ (from Robert J Johnston Jr) is a null allele for both *BEAF-32A* and *BEAF-32B*. *CP190[1]* and *CP190[2]* ^92^ were obtained from Jordan Raff. Live muscle twitching movies were recorded by labelling the muscle ends with Talin-C-terminal-YPet ^93^.

#### Generation of tono[1] and tono[2] alleles

*tono[1]* was generated as described with the published TALEN method ^45^. Briefly, two pairs of TALENs (pair 1 targeting AACCGCAGCATCATC in the first coding exon of *tono* and pair 2 targeting ACTCCGGAGAGGTGA in the second coding exon of *tono*) have been designed and constructed ^94^. *In vitro* transcribed TALEN mRNAs were injected into *w*^1118^ fly embryos at a concentration of 250 ng/µl each. The genomic region covering both targeted sites has been amplified by PCR with primer XZ13 and XZ14 and then followed by a T7 assay to identify successful targeting ^45^. Fly stocks with mutations in *tono* were established and sequenced.

*tono[2]* was constructed by CRPSPR-RMCE genome editing as published with small optimisations ^46^. In brief, the intron sequence between coding exons 1 and 2 of *tono* and sequence from the end of 3’-UTR of *tono* were chosen for sgRNA designing with CRISPRscan ^95^. Four sgRNAs were picked for each targeting region according to the score from CRISPRscan and their activities were estimated in S2 cells as described^46^. sgRNA 1 (targeting sequences GTGTAATGCGGTGAAAGCG in the intron region) and sgRNA 2 (targeting sequences GGGTCTAAGACGTTGGTTT in 3’-UTR) were picked for embryo injection. The left homology arm of ∼2kb and the right arm of ∼2kb were assembled with dsRed cassette as a repair donor as described ^46^. The *y[1], M(Act5C-Cas9)ZH-2A, w[1118], DNAlig4[169]* embryos were injected with donor plasmid at 800 ng/µl and sgRNAs at 272 ng/µl each. F1 flies with red fluorescent eyes were chosen for characterisation by PCR and sequencing to confirm the cassette insertion. Fly stocks with correct insertion were established as *tono[2]*. For the RMCE exchange, a full length of *tono* coding sequences (CDS), *tono-ΔBTB*, and *tono-ΔZFs* were tagged with 2xHA and cloned as described ^46^. The plasmids were injected into *tono[2]* fly embryos and the progeny were screened for non-fluorescent eyes and characterised by PCR and sequencing as described ^46^.

#### Transcriptomics

For transcriptomics, flight muscles from wildtype control and *tono[1]* flies were dissected at either 30 h APF or 48 h APF and isolated based on *Mef2*-GAL4, UAS-GFP-Gma expression ^24^. Three replicates were used and each replicate contained indirect flight muscles dissected from 150 pupae. mRNA libraries were prepared as described previously ^24^. Libraries were sequenced on an Illumina HiSeq 2500 and multiplexed 3-4 samples per lane, obtaining ∼100 million reads per library. Sequencing was performed at the Vienna Biocenter Core Facility (https://www.viennabiocenter.org/vbcf). Reads were filtered and trimmed using the FASTX Toolkit and cutadapt then mapped to the Flybase 2015_04 genome assembly using STAR {Dobin:2013fg}. Reads were visualized on the USCS server by normalizing them to the largest library size. featureCounts v1.4.2 ^96^ was used for determining raw read counts. Differential expression analysis was done using the RNfuzzyApp ^97^, with the TCC normalization ^98^ and DESeq2 ^99^ method. Additional data processing was handled in R.

#### Phasik analysis

Phasik was used for mitochondrial phase detection. We used the mito-interactome from mitoXplorer 2.0 ^52^ as network, together with the transcriptomic data from the flight muscle developmental time-course ^19^ to construct the temporal network. We used Euclidian distance and hierarchical clustering with the ‘Ward’ method for temporal network clustering. As Phasik can detect at multiple scales, we chose to analyse the three main clusters, which reflect the prominent switch between 30 h and 48 h APF in the distance matrix, and a second switch between 72 h and 90 h AFP. An edge weight of 0.7 was chosen to select significant genes for enrichment analysis, which was then done with FlyEnrichR ^100^.

#### Immunostaining and processing

Flight muscle dissections were performed with a slightly modified published protocol^101^. Briefly, after removing the pupal case from the staged pupae, two to three holes were pinned in the abdomen with insect pins. The samples were then transferred to 4% PFA in PBST (0.5% Triton-X in PBS) and fixed for 20 min at room temperature (RT) on a rocking platform. Samples were transferred to a silicon-coated petri dish and their position was fixed as described ^101^. One hole was cut from the head to the abdomen along the ventral middle line. Other tissues in between the two thorax halves were removed by forceps or by pipetting. The two thorax halves were then cut off and transferred to a 24-well plate. The samples were washed with 3x PBST for 10 min each at RT before blocking with 5% normal goat serum (NGS) in PBST for 1 hour at RT. Afterwards, they were incubated with primary antibodies at 4°C overnight and washed 3x in PBST (10 min each) at RT. Incubation with secondary antibodies was performed at room temperature for 2 hours. The samples were washed 3x in PBST and mounted in VECTASHIELD plus DAPI (Vector) to visualise nuclei. The samples were stored at 4°C before imaging. Images were acquired on Zeiss LSM780 or LSM880 confocal with AiryScan and processed with Fiji (ImageJ).

To stain for Su(Hw) and Mod(mdg4)67.2 the samples were dissected and then incubated for 30 min in PBS or PBS + 250mM NaCl before fixing with methanol. The fixation was done in precooled 100% methanol at −20 °C for 15 min with gentle rocking every 5 min. Afterwards, an equal volume of PBST was added and samples were washed as described above before proceeding to immunostaining.

Rabbit anti-GFP (1:2000) was obtained from Amsbio (TP401). Rat monoclonal anti-HA (3F10) and mouse monoclonal anti-HA (16B12) were used at 1:500. Rabbit anti-CP190 (dilution 1:2000), mouse anti-CP190 (dilution 1:1000), rabbit anti-Mod(mdg4)67.2 (dilution 1:1000) and rabbit anti-Su(Hw) (1:1000) were gifts from Mariano Labrador ^54^. Mouse anti-BEAF-32 (dilution 1:20) ^102^, mouse anti-βPS-integrin (CF.6G11, dilution 1:100) ^103^, mouse anti-Futsch (22C10, dilution 1:100) ^104^, mouse anti-LaminDm0 (ADL67.10, dilution 1:50) ^105^, and mouse anti-HP1a (C19A, 1:100) ^106^ were obtained from DSHB. Rabbit anti-Gelsolin (dilution, 1:2000) ^107^ was a gift from Maria Leptin. All the primary antibodies were diluted in PBST with 5% NGS. All the secondary antibodies, including goat anti-mouse, goat anti-rat, goat anti-pig, and goat anti-rabbit, and phalloidin-rhodamine were purchased from Molecular Probes and applied at 1:500 dilution. Tono-GFP signal was boosted after fixation with a GFP nanobody (gift of Dirk Görlich) at 1:2000 for 2 hours at room temperature.

Sarcomere length quantification: Fly pupal samples were staged and dissected as described ^101^. The flight muscles were stained with rhodamine phalloidin and images were acquired with a Leica LSM780 or LSM880 confocal. Sarcomere length was measured using the Fiji plug-in MyofibrilJ ^19^. Two-tailed unpaired student t-tests were performed between wild type and *tono[1]* or *tono[2]*. Error bar represents the standard deviation.

Mitochondria were labelled by expressing mitochondrial matrix-directed GFP (UAS-mitGFP, BDSC: 25747). Quantification of the aspect ratio (length / width) of the mitochondria was performed in double-blind fashion using hand segmentation, systematically measuring the long and short axes of each mitochondrion from a cropped image of 40×20µm. Mitochondria were quantified from n>4 thoraxes. The total area of mitochondria is identified by Otsu thresholding on Fiji for each z-plane. The multiple quantifications for each z-plane were averaged for each animal (one value per animal)^34^.

Tono-granularity: To quantify the Tono granularity score, each nucleus was segmented in 2D in one z-plane. For each nucleus (ROI) the mean signal and its standard deviation was retrieved to quantify the granularity score (mean/std) using a Fiji macro (https://github.com/PierreMangeol/Tono). To verify our methodology, we devised a simple simulation in Python to mimic the different images of Tono condensation states we observed. We then quantified these images the same way we analysed Tono distributions in the muscle nuclei. In this simulation, the total amount of Tono is constant, and at a given state of condensation, Tono condensates are spheres of equal radii that cannot overlap each other. Spheres spread over a two-dimensional square area that is the same for the entire simulation. To simulate images of these condensates, images of these spheres were simplified as discs and to take into account the diffraction-limited nature of microscopy observations, images were convoluted with a Gaussian kernel. The resulting simulated images are comparable to real microscopy images that would be obtained with a resolution of 210 nm (wavelength of 550 nm observed with an objective of N.A. 1.3) and particle diameters ranging from 100 nm to 700 nm. This simulation is an oversimplification of reality, but it recapitulates well the fact that the mean-normalised standard deviation of intensity of Tono in nuclei increases together with the condensation process (Supplemental Figure 8).

#### Osmotic and mechanical manipulations

To induce hyper-osmotic stress, pupae were taken out of the pupal case and pierced 3 times in the abdomen with a dissecting needle and then incubated in PBS control or PBS + 250 mM NaCl or PBS + 500 mM sucrose at RT for 30 min on a belly dancer before fixation. The reversibility of the Tono droplets was tested by continuing with a hypo-osmotic condition by further incubating the samples in 0.1x PBS for 20 min or keeping them in PBS control buffer.

Mechanical manipulations were performed by preparing living adult hemi-thoraxes from Tono-GFP flies, without damaging the flight muscles ^34^ and mounting them between slides and a coverslip using either 4 or 2 layers of Scotch tape on each side as spacer for the relaxed and squeezed conditions, respectively. The flight muscles were imaged for the next 20 minutes after the dissection.

#### Life imaging

To observe the dynamics of Tono-GFP, living intact living flight muscles were mounted as described above and imaged with a spinning disc confocal. To image muscle twitching, we used Talin-C-terminal-YPet and followed published protocols ^19,108^.

#### Flight test

Flight tests were performed as previously described ^93^. Around 20-30 fresh adult males (1-3 days after eclosion) were collected and recovered for 48 hours in the incubator at 27 degrees. Flies were introduced into a flight test cylinder divided into 3 zones and the landing position of the flies was documented.

## Acknowledgements

We thank the Schnorrer lab and Dirk Görlich for insightful discussions and input on this manuscript. We thank Johannes Bischof, Dirk Görlich, Robert J Johnston Jr, Mariano Labrador, Maria Leptin, Jordan Raff and Rainer Renkawitz for generous gifts of antibodies and fly strains. We are grateful to the DSHB for antibodies and the Bloomington *Drosophila* Stock Center for fly strains (NIH P40OD018537) and to Flybase for database organisation. We are grateful to Christophe Pitaval for fly embryo injections and fly stock maintenance. We thank Fabio Marchiano for generous help in bioinformatics. We are indebted to the IBDM imaging and fly facilities for help with image acquisition, maintenance of the microscopes and fly food.

## Supplementary legends

**Supplemental Figure 1: *tono* mutant phenotype is muscle autonomous**

**(A)** *Drosophila* adult flight muscle schemes. Scheme of an adult fly with dorsal longitudinal flight muscles in magenta (left). Scheme of the adult thorax (middle) and one individual muscle fiber (right) (**B-J**): Adult hemi-thoraces of heterozygous *tono* deficiency *Df(3L)ED4502* (B), *tono[1] / Df(3L)ED4502* (C), *tono[2] / Df(3L)ED4502* (D), *tono-GFP* tagged Fosmid (E), *tono[1], tono-GFP* (F), *Mhc[10]; tono[1]* (G), *Act88F-GAL4*; *tono[1]* (H), *Act88F-GAL4*; *tono[1], UAS-tono-HA* (I) and *Act88F-GAL4*; *tono[2], UAS-tono-HA* (J) stained with rhodamine phalloidin to label F-actin. Note that muscle-specific expression of *tono* rescues the muscle atrophy. Scale bar represents 100 µm.

**Supplemental Figure 2: *tono* mutant developmental phenotype**

**(A)** Schemes of 56 h APF wild-type (left) and *tono[1]* mutant thoraces (right), dorsal longitudinal flight muscles are in magenta, tendons in green and cuticle in black. (**B-M**) Developing flight muscles from wild type (B-E), *tono[1]* (F-I) and *tono[2]* (J-M) of the indicated time points stained with phalloidin. Note the muscle detachments indicated with the red arrowheads and the fiber ruptures indicated with the green arrowheads. (**N-P**) Flight muscle motor neurons stained anti-Futsch in green of wild-type (N), *tono[1]* (O) and *tono[2]* (P) mutant 48 h APF pupae. Scale bars represent 100 µm.

**Supplemental Figure 3: *tono* expression and domain function**

**(A)** *tono* (*CG32121*) genomic map visualised with the UCSC genome browser. Magenta boxes are predicted coding exons, and UTR regions are in yellow. Below are mRNA read counts from published developmental transcriptomics data ^19^. Note the high mRNA expression at 30 h and 48 h APF. (B-G). Flight muscles (B-G) and sarcomeres (H-J) visualised by phalloidin staining of *tono-HA, tono-ΔBTB-HA* and *tono-ΔZfs-HA* alleles. Scale bars represent 100 µm in B-G and 5 µm in H-J.

**Supplemental Figure 4: Tono regulates mitochondrial gene expression**

(**A-B**): Volcano plots of of *tono[1]* vs. wild type RNA-Seq at 30 h APF (A) and 48 h APF (B). Significantly changed genes (DESeq2, FDR<0.05, log2FC>1) are in red, and all other genes are in green. All data are listed in Supplemental data 1. (**C**) Changes in expression levels of different mitochondrial process genes in wild type or *tono[1]* 48 h vs 30 h APF visualised by the mitoXplorer suite ^52^. Note that genes down-regulated (red) in wild type are less down-regulated in *tono[1]*, genes up-regulated (blue) in wild type are less up-regulated in *tono[1]*. Key glycolysis (Pfk) and pyruvate metabolism (LDH) enzymes are highlighted in red.

**Supplemental Figure 5: *tono* regulates a transcriptional switch at 30 h to 48 h APF**

Violin plots comparing expression of 48 h vs 30 h APF from genes contained in each of the 40 developmental mRNA clusters identified during wild-type flight muscle development in ^19^. Note that many clusters up-regulated in wild type are less up-regulated in *tono[2]* (blue), and clusters that are down-regulated in wild type are less down-regulated in *tono[2]* (red).

**Supplemental Figure 6: Tono forms liquid droplets upon hyperosmotic treatment**

(**A-D**) Tono-HA localisation pattern in flight muscles at 72 h APF or one-day adult either incubated with isotonic (A, B) or hypertonic 250 mM NaCl (C, D) solution for 30 min. anti-HA is shown in green and DAPI in magenta. Note the large phase-separated Tono droplets after hypertonic treatment. (**E**) Tono-GFP droplet formation in 48 h APF flight muscles after incubation with 500 mM sucrose solution for 30 min. (**F, G**) Control (F) and hypertonic (250 mM NaCl, G) incubation of 48 h APF Tono-GFP flight muscles stained with anti-HP1 in red and anti-GFP in green. All scale bars represent 5 µm.

**Supplemental Figure 7: Tono droplets are distinct from insulator bodies**

(**A-C**) Control (A) and hypertonic (250 mM NaCl, B, C) incubation of 48 h APF Tono-GFP (A, B) or wild-type (C) flight muscles stained with anti-GFP in green and anti-BEAF-32 in red (A, B), or anti-BEAF-32 in green and anti-CP190 in red (C). (**D-G**) Control (D, E) and hypertonic (250 mM NaCl, F, G) incubation of 48 h APF Tono-HA flight muscles stained with anti-HA in green and anti-Su(Hw) in red (D, F) or anti-Mod(mdg4)67.2 in red (E, G). (**H-K**) Flight muscles from pharate or 1-day adults of *CTCF^GE^*^24185^ (H), *CTCF^GE^*^24185^*^/P^*^30^*^.6^* (I), *BEAF-32^[AB-KO]^* (J), and *CP190^1/2^* (K). Scale bars represent 5 µm in A-G and 100 µm in H-K.

**Supplemental Figure 8: Tono granularity simulation**

Simulated image of Tono particles of increasing size, simulating the condensation process of Tono in nuclei (lower images). The simulated images are comparable to real microscopy images that would be obtained with a resolution of 210 nm (wavelength of 550 nm observed with an objective of N.A. 1.3) and particle diameters ranging from 100 nm to 700 nm. Mean-normalised standard deviation of simulated images plotted against particle radius (top). Note the increase in the mean-normalized standard deviation with increasing particle size.

**Supplemental Movie 1: Muscle twitching at 48 h APF**

Live movies of developing wild-type (left) or *tono[2]* (right) mutant flight muscles at 48 h APF that were labelled with Talin-C-YPet. Scale bar represents 50 µm.

**Supplemental Movie 2: Muscle twitching at 60 h APF**

Live movies of developing wild-type (left) or *tono[2]* (right) mutant flight muscles at 60 h APF that were labelled with Talin-C-YPet. Note the twitch in the *tono[2]* muscles. Scale bar represents 50 µm.

**Supplemental Movie 3: Tono-GFP phase separation induced by mechanical pressure**

Live movie of Tono-GFP flight muscles that were mechanically squeezed. Not the fast Tono-GFP droplet formation in the nuclei. Scale bar represents 5 µm.

**Supplemental Movie 4: Tono-GFP droplets fuse**

High-speed live movie of Tono-GFP in one flight muscle nucleus that was mechanically squeezed. Not the fast Tono-GFP droplets fuse. The stills in Figure 6N correspond to frames starting at 50 sec of this movie. Scale bar represents 5 µm.

**Supplemental Data 1. Raw, normalised and differential mRNA-Seq data, enrichment analysis**

The file includes multiple tabs containing the mRNA-Seq raw counts data, TCC-DESeq2 normalized and differential expression data either showing all genes, or all significantly changed genes (log2FC <-1, >1 and FDR <0.05). Comparison of wild type vs. *tono*[1] at 30 h APF and 48 h APF are listed (metabolic gene are highlighted in yellow and cytoskeletal ones in blue, innervation ones in red and muscle transcription factors in grey), as well as 30 h vs. 48 h APF wild type and 30 h vs. 48 APF *tono*[1]. The last tab displays the fly-EnrichR analysis of 48 h vs. *tono[1]* listing KEGG pathway and GO-term Cellular Component; metabolic pathways are highlighted in yellow and cytoskeletal terms in blue.

**Supplemental Data 2. Phasik result data**

The file displays the results Phasik ^53^ using the mitochondrial interactome ^52^ and the RNA-seq data from flight muscle development mRNA-Seq data ^19^. We chose to analyse 3 phases: phase 1: myoblast to 30 h APF, phase 2: 48 h to 72 h APF and phase 3: 90 h APF to adult stage. The file includes multiple tabs, with the interacting proteins (i, j) of each cluster together with the edge weight. KEGG enrichment in FlyEnrichR () was done with the most prominent interacting partners, with weight > 0.7.

## References

1. Ehler, E. & Gautel, M. The sarcomere and sarcomerogenesis. Adv. Exp. Med. Biol. 642, 1–14 (2008).

2. Lange, S., Ehler, E. & Gautel, M. From A to Z and back? Multicompartment proteins in the sarcomere. Trends in Cell Biology 16, 11–18 (2006).

3. Loison, O. et al. Polarization-resolved microscopy reveals a muscle myosin motor-independent mechanism of molecular actin ordering during sarcomere maturation. PLoS Biol 16, e2004718 (2018).

4. Dasbiswas, K., Hu, S., Schnorrer, F., Safran, S. A. & Bershadsky, A. D. Ordering of myosin II filaments driven by mechanical forces: experiments and theory. Philos. Trans. R. Soc. Lond., B, Biol. Sci. 373, 20170114 (2018).

5. Schiaffino, S. & Reggiani, C. Fiber types in mammalian skeletal muscles. Physiological Reviews 91, 1447–1531 (2011).

6. Spletter, M. L. & Schnorrer, F. Transcriptional regulation and alternative splicing cooperate in muscle fiber-type specification in flies and mammals. Experimental Cell Research 321, 90–98 (2014).

7. Schönbauer, C. et al. Spalt mediates an evolutionarily conserved switch to fibrillar muscle fate in insects. Nature 479, 406–409 (2011).

8. Vigoreaux, J. O. in Muscle Development in Drosophila 69, 143–156 (Springer New York, 2006).

9. Lemke, S. B. & Schnorrer, F. Mechanical forces during muscle development. Mechanisms of Development 144, 92–101 (2017).

10. Kim, J. H., Jin, P., Duan, R. & Chen, E. H. Mechanisms of myoblast fusion during muscle development. Current Opinion in Genetics & Development 32, 162–170 (2015).

11. Schweitzer, R., Zelzer, E. & Volk, T. Connecting muscles to tendons: tendons and musculoskeletal development in flies and vertebrates. Development 137, 2807–2817 (2010).

12. Schnorrer, F. & Dickson, B. J. Muscle building; mechanisms of myotube guidance and attachment site selection. 7, 9–20 (2004).

13. Sanger, J. W., Wang, J., Fan, Y., White, J. & Sanger, J. M. Assembly and dynamics of myofibrils. Journal of Biomedicine and Biotechnology 2010, 858606 (2010).

14. Sparrow, J. C. & Schöck, F. The initial steps of myofibril assembly: integrins pave the way. Nature Reviews Molecular Cell Biology 10, 293–298 (2009).

15. Weitkunat, M., Kaya-Copur, A., Grill, S. W. & Schnorrer, F. Tension and force-resistant attachment are essential for myofibrillogenesis in Drosophila flight muscle. Curr Biol 24, 705–716 (2014).

16. Weitkunat, M., Brasse, M., Bausch, A. R. & Schnorrer, F. Mechanical tension and spontaneous muscle twitching precede the formation of cross-striated muscle in vivo. Development 144, 1261–1272 (2017).

17. Sanger, J. W. et al. Assembly and Maintenance of Myofibrils in Striated Muscle. Handb Exp Pharmacol 235, 39–75 (2017).

18. Sanger, J. W., Wang, J., Holloway, B., Du, A. & Sanger, J. M. Myofibrillogenesis in skeletal muscle cells in zebrafish. Cell Motil. Cytoskeleton 66, 556–566 (2009).

19. Spletter, M. L. et al. A transcriptomics resource reveals a transcriptional transition during ordered sarcomere morphogenesis in flight muscle. eLife 7, 1361 (2018).

20. Orfanos, Z. et al. Sallimus and the dynamics of sarcomere assembly in Drosophila flight muscles. Journal of Molecular Biology 427, 2151–2158 (2015).

21. Johnston, J. J. et al. A novel nemaline myopathy in the Amish caused by a mutation in troponin T1. Am. J. Hum. Genet. 67, 814–821 (2000).

22. Robinson, P. et al. Mutations in fast skeletal troponin I, troponin T, and beta-tropomyosin that cause distal arthrogryposis all increase contractile function. FASEB J. 21, 896–905 (2007).

23. Patel, N. et al. ZBTB42 mutation defines a novel lethal congenital contracture syndrome (LCCS6). Human Molecular Genetics 23, 6584–6593 (2014).

24. Spletter, M. L. et al. The RNA-binding protein Arrest (Bruno) regulates alternative splicing to enable myofibril maturation in Drosophila flight muscle. EMBO Rep 16, 178–191 (2015).

25. Nongthomba, U., Cummins, M., Clark, S., Vigoreaux, J. O. & Sparrow, J. C. Suppression of muscle hypercontraction by mutations in the myosin heavy chain gene of Drosophila melanogaster. 164, 209–222 (2003).

26. Reedy, M., Bullard, B. & Vigoreaux, J. Flightin is essential for thick filament assembly and sarcomere stability in Drosophila flight muscles. Journal of Cell Biology 151, 1483 (2000).

27. Firdaus, H. et al. A cis-regulatory mutation in troponin-I of Drosophila reveals the importance of proper stoichiometry of structural proteins during muscle assembly. 200, 149–165 (2015).

28. Burden, S. J., Huijbers, M. G. & Remedio, L. Fundamental Molecules and Mechanisms for Forming and Maintaining Neuromuscular Synapses. Int J Mol Sci **19**, (2018).

29. Elhanany-Tamir, H. et al. Organelle positioning in muscles requires cooperation between two KASH proteins and microtubules. The Journal of Cell Biology 198, 833–846 (2012).

30. Roman, W. & Gomes, E. R. Nuclear positioning in skeletal muscle. Semin. Cell Dev. Biol. 82, 51–56 (2018).

31. Cornelissen, T. et al. Deficiency of parkin and PINK1 impairs age-dependent mitophagy in Drosophila. eLife 7, 487 (2018).

32. Park, J. et al. Mitochondrial dysfunction in Drosophila PINK1 mutants is complemented by parkin. Nature 441, 1157–1161 (2006).

33. Sauerwald, J., Backer, W., Matzat, T., Schnorrer, F. & Luschnig, S. Matrix metalloproteinase 1 modulates invasive behavior of tracheal branches during entry into Drosophila flight muscles. eLife 8, 1079 (2019).

34. Avellaneda, J. et al. Myofibril and mitochondria morphogenesis are coordinated by a mechanical feedback mechanism in muscle. Nature Communications 12, 2091–18 (2021).

35. Lin, J. et al. Transcriptional co-activator PGC-1 alpha drives the formation of slow-twitch muscle fibres. Nature 418, 797–801 (2002).

36. Poliacikova, G. et al. M1BP is an essential transcriptional activator of oxidative metabolism during Drosophila development. Nature Communications 14, 3187–20 (2023).

37. Weitkunat, M., Kaya-Copur, A., Grill, S. W. & Schnorrer, F. Tension and force-resistant attachment are essential for myofibrillogenesis in Drosophila flight muscle. Curr Biol 24, 705–716 (2014).

38. Bernstein, S. I., Mogami, K., Donady, J. J. & Emerson, C. P. Drosophila muscle myosin heavy chain encoded by a single gene in a cluster of muscle mutations. Nature 302, 393–397 (1983).

39. Beall, C. J., Sepanski, M. A. & Fyrberg, E. A. Genetic dissection of Drosophila myofibril formation: effects of actin and myosin heavy chain null alleles. Genes & Development 3, 131–140 (1989).

40. Cripps, R. M., Ball, E., Stark, M., Lawn, A. & Sparrow, J. C. Recovery of dominant, autosomal flightless mutants of Drosophila melanogaster and identification of a new gene required for normal muscle structure and function. 137, 151–164 (1994).

41. van Dijk, S. J. et al. Cardiac myosin-binding protein C mutations and hypertrophic cardiomyopathy: haploinsufficiency, deranged phosphorylation, and cardiomyocyte dysfunction. Circulation 119, 1473–1483 (2009).

42. Marston, S. et al. Evidence from human myectomy samples that MYBPC3 mutations cause hypertrophic cardiomyopathy through haploinsufficiency. Circ. Res. 105, 219–222 (2009).

43. Schnorrer, F. et al. Systematic genetic analysis of muscle morphogenesis and function in Drosophila. Nature 464, 287–291 (2010).

44. Li, H. et al. Fly Cell Atlas: A single-nucleus transcriptomic atlas of the adult fruit fly. 375, eabk2432 (2022).

45. Zhang, X., Ferreira, I. R. S. & Schnorrer, F. A simple TALEN-based protocol for efficient genome-editing in Drosophila. METHODS 69, 32–37 (2014).

46. Zhang, X., Koolhaas, W. H. & Schnorrer, F. A versatile two-step CRISPR-and RMCE-based strategy for efficient genome engineering in Drosophila. G3 (Bethesda) 4, 2409–2418 (2014).

47. Sarov, M. et al. A genome-wide resource for the analysis of protein localisation in Drosophila. eLife 5, e12068 (2016).

48. Cripps, R. M., Suggs, J. A. & Bernstein, S. I. Assembly of thick filaments and myofibrils occurs in the absence of the myosin head. The EMBO Journal 18, 1793–1804 (1999).

49. Venken, K. J. T. et al. MiMIC: a highly versatile transposon insertion resource for engineering Drosophila melanogaster genes. Nature Methods 8, 737–743 (2011).

50. Kanca, O., Bellen, H. J. & Schnorrer, F. Gene Tagging Strategies To Assess Protein Expression, Localization, and Function in Drosophila. 207, 389–412 (2017).

51. Siggs, O. M. & Beutler, B. The BTB-ZF transcription factors. Cell Cycle 11, 3358–3369 (2012).

52. Marchiano, F., Haering, M. & Habermann, B. H. The mitoXplorer 2.0 update: integrating and interpreting mitochondrial expression dynamics within a cellular context. Nucleic Acids Res. (2022). doi:10.1093/nar/gkac306

53. Lucas, M. et al. Inferring cell cycle phases from a partially temporal network of protein interactions. Cell Reports Methods 100397 (2023). doi:10.1016/j.crmeth.2023.100397

54. Schoborg, T., Rickels, R., Barrios, J. & Labrador, M. Chromatin insulator bodies are nuclear structures that form in response to osmotic stress and cell death. The Journal of Cell Biology 202, 261–276 (2013).

55. Strom, A. R. et al. Phase separation drives heterochromatin domain formation. Nature 547, 241–245 (2017).

56. Larson, A. G. et al. Liquid droplet formation by HP1α suggests a role for phase separation in heterochromatin. Nature 547, 236–240 (2017).

57. Mohan, M. et al. The Drosophila insulator proteins CTCF and CP190 link enhancer blocking to body patterning. The EMBO Journal 26, 4203–4214 (2007).

58. Cavalheiro, G. R. et al. CTCF, BEAF-32, and CP190 are not required for the establishment of TADs in early Drosophila embryos but have locus-specific roles. Sci Adv 9, eade1085 (2023).

59. Gambetta, M. C. & Furlong, E. E. M. The Insulator Protein CTCF Is Required for Correct Hox Gene Expression, but Not for Embryonic Development in Drosophila. (2018). doi:10.1534/genetics.118.301350

60. Kaushal, A. et al. CTCF loss has limited effects on global genome architecture in Drosophila despite critical regulatory functions. Nature Communications 1–16 (2021). doi:10.1038/s41467-021-21366-2

61. Kyrchanova, O. et al. Drosophila architectural protein CTCF is not essential for fly survival and is able to function independently of CP190. Biochim Biophys Acta Gene Regul Mech 1864, 194733 (2021).

62. Frey, S., Richter, R. P. & Görlich, D. FG-rich repeats of nuclear pore proteins form a three-dimensional meshwork with hydrogel-like properties. Science 314, 815–817 (2006).

63. Shin, Y. & Brangwynne, C. P. Liquid phase condensation in cell physiology and disease. Science 357, (2017).

64. Hyman, A. A., Weber, C. A. & JUlicher, F. Liquid-liquid phase separation in biology. 30, 39–58 (2014).

65. Alberti, S. & Hyman, A. A. Biomolecular condensates at the nexus of cellular stress, protein aggregation disease and ageing. Nature Reviews Molecular Cell Biology 22, 196–213 (2021).

66. Nordgaard, C. et al. ZAKβ is activated by cellular compression and mediates contraction-induced MAP kinase signaling in skeletal muscle. The EMBO Journal e111650 (2022). doi:10.15252/embj.2022111650

67. Venkova, L. et al. A mechano-osmotic feedback couples cell volume to the rate of cell deformation. eLife 11, (2022).

68. Luis, N. M. & Schnorrer, F. Mechanobiology of muscle and myofibril morphogenesis. Cells & Development 168, 203760 (2021).

69. Swist, S. et al. Maintenance of sarcomeric integrity in adult muscle cells crucially depends on Z-disc anchored titin. Nature Communications 11, 4479–18 (2020).

70. Roman, W. et al. Myofibril contraction and crosslinking drive nuclear movement to the periphery of skeletal muscle. Nature cell biology 19, 1189– 1201 (2017).

71. Barbas, J. A., Galceran, J., Torroja, L., Prado, A. & Ferrús, A. Abnormal muscle development in the heldup3 mutant of Drosophila melanogaster is caused by a splicing defect affecting selected troponin I isoforms. Molecular and Cellular Biology 13, 1433–1439 (1993).

72. Nongthomba, U., Ansari, M., Thimmaiya, D., Stark, M. & Sparrow, J. Aberrant splicing of an alternative exon in the Drosophila troponin-T gene affects flight muscle development. 177, 295–306 (2007).

73. Nongthomba, U. et al. Troponin I is required for myofibrillogenesis and sarcomere formation in Drosophila flight muscle. 117, 1795–1805 (2004).

74. Fischbarg, J. Fluid transport across leaky epithelia: central role of the tight junction and supporting role of aquaporins. Physiological Reviews 90, 1271– 1290 (2010).

75. Torres-Sánchez, A., Kerr Winter, M. & Salbreux, G. Tissue hydraulics: Physics of lumen formation and interaction. Cells & Development 168, 203724 (2021).

76. Lafontaine, D. L. J., Riback, J. A., Bascetin, R. & Brangwynne, C. P. The nucleolus as a multiphase liquid condensate. Nature Reviews Molecular Cell Biology 1–18 (2021). doi:10.1038/s41580-020-0272-6

77. Amankwaa, B., Schoborg, T. & Labrador, M. Drosophila insulator proteins exhibit in vivo liquid-liquid phase separation properties. Life Sci. Alliance 5, (2022).

78. Hnisz, D., Shrinivas, K., Young, R. A., Chakraborty, A. K. & Sharp, P. A. A Phase Separation Model for Transcriptional Control. CELL 169, 13–23 (2017).

79. Dupont, S. & Wickström, S. A. Mechanical regulation of chromatin and transcription. Nat Rev Genet 23, 624–643 (2022).

80. Goswami, R. et al. Mechanical Shielding in Plant Nuclei. Curr Biol 30, 2013–2025.e3 (2020).

81. Venturini, V. et al. The nucleus measures shape changes for cellular proprioception to control dynamic cell behavior. 370, (2020).

82. Lomakin, A. J. et al. The nucleus acts as a ruler tailoring cell responses to spatial constraints. Science 370, (2020).

83. Swift, J. et al. Nuclear lamin-A scales with tissue stiffness and enhances matrix-directed differentiation. 341, 1240104 (2013).

84. Wang, S. et al. Mechanotransduction via the LINC complex regulates DNA replication in myonuclei. The Journal of Cell Biology 217, 2005–2018 (2018).

85. Heller, S. A., Shih, R., Kalra, R. & Kang, P. B. Emery-Dreifuss muscular dystrophy. Muscle Nerve **61**, 436–448 (2020).

86. Patel, N. et al. ZBTB42 mutation defines a novel lethal congenital contracture syndrome (LCCS6). Human Molecular Genetics 23, 6584–6593 (2014).

87. Bischof, J. et al. A versatile platform for creating a comprehensive UAS-ORFeome library in Drosophila. Development 140, 2434–2442 (2013).

88. Bryantsev, A. L., Baker, P. W., Lovato, T. L., Jaramillo, M. S. & Cripps, R. M. Differential requirements for Myocyte Enhancer Factor-2 during adult myogenesis in Drosophila. Dev. Biol. 361, 191–207 (2012).

89. Collier, V. L., Kronert, W. A., O’Donnell, P. T., Edwards, K. A. & Bernstein, S. I. Alternative myosin hinge regions are utilized in a tissue-specific fashion that correlates with muscle contraction speed. Genes & Development 4, 885– 895 (1990).

90. Fernandes, J., Celniker, S. & VijayRaghavan, K. Development of the Indirect Flight Muscle Attachment Sites inDrosophila: Role of the PS Integrins and thestripeGene. 176, 166–184 (1996).

91. Roy, S., Gilbert, M. K. & Hart, C. M. Characterization of BEAF mutations isolated by homologous recombination in Drosophila. Genetics 176, 801–813 (2007).

92. Butcher, R. D. J. et al. The Drosophila centrosome-associated protein CP190 is essential for viability but not for cell division. 117, 1191–1199 (2004).

93. Lemke, S. B., Weidemann, T., Cost, A.-L., Grashoff, C. & Schnorrer, F. A small proportion of Talin molecules transmit forces at developing muscle attachments in vivo. PLoS Biol 17, e3000057 (2019).

94. Cermak, T., et al. Efficient design and assembly of custom TALEN and other TAL effector-based constructs for DNA targeting. 39, e82–e82 (2011).

95. Moreno-Mateos, M. A. et al. CRISPRscan: designing highly efficient sgRNAs for CRISPR-Cas9 targeting in vivo. Nature Methods 12, 982–988 (2015).

96. Liao, Y., Smyth, G. K. & Shi, W. featureCounts: an efficient general purpose program for assigning sequence reads to genomic features. Bioinformatics 30, 923–930 (2014).

97. Haering, M. & Habermann, B. H. RNfuzzyApp: an R shiny RNA-seq data analysis app for visualisation, differential expression analysis, time-series clustering and enrichment analysis. F1000Research 2021 10:654 **10**, 654 (2021).

98. Sun, J., Nishiyama, T., Shimizu, K. & Kadota, K. TCC: Differential expression analysis for tag count data with robust normalization strategies. (2023).

99. Love, M. I., Huber, W. & Anders, S. Moderated estimation of fold change and dispersion for RNA-seq data with DESeq2. Genome Biology 15, 550 (2014).

100. Kuleshov, M. V., et al. modEnrichr: a suite of gene set enrichment analysis tools for model organisms. 47, W183–W190 (2019).

101. Weitkunat, M. & Schnorrer, F. A guide to study Drosophila muscle biology. METHODS 68, 2–14 (2014).

102. Blanton, J., Gaszner, M. & Schedl, P. Protein:protein interactions and the pairing of boundary elements in vivo. Genes & Development 17, 664–675 (2003).

103. Brower, D. L., Wilcox, M., Piovant, M., Smith, R. J. & Reger, L. A. Related cell-surface antigens expressed with positional specificity in Drosophila imaginal discs. Proceedings of the National Academy of Sciences of the United States of America 81, 7485–7489 (1984).

104. Zipursky, S. L., Venkatesh, T. R., Teplow, D. B. & Benzer, S. Neuronal development in the Drosophila retina: monoclonal antibodies as molecular probes. CELL 36, 15–26 (1984).

105. Riemer, D. et al. Expression of Drosophila lamin C is developmentally regulated: analogies with vertebrate A-type lamins. J Cell Sci 108 **(Pt** **10**), 3189–3198 (1995).

106. James, T. C. & Elgin, S. C. Identification of a nonhistone chromosomal protein associated with heterochromatin in Drosophila melanogaster and its gene. Molecular and Cellular Biology 6, 3862–3872 (1986).

107. Stella, M. C., Schauerte, H., Straub, K. L. & Leptin, M. Identification of secreted and cytosolic gelsolin in Drosophila. Journal of Cell Biology 125, 607–616 (1994).

108. Lemke, S. B. & Schnorrer, F. In Vivo Imaging of Muscle-tendon Morphogenesis in Drosophila Pupae. JoVE e57312–e57312 (2018). doi:10.3791/57312

