## Supplementary Figures 1-8 for "A mechanically regulated liquid-liquid phase separation of the transcriptional regulator Tono instructs muscle development"

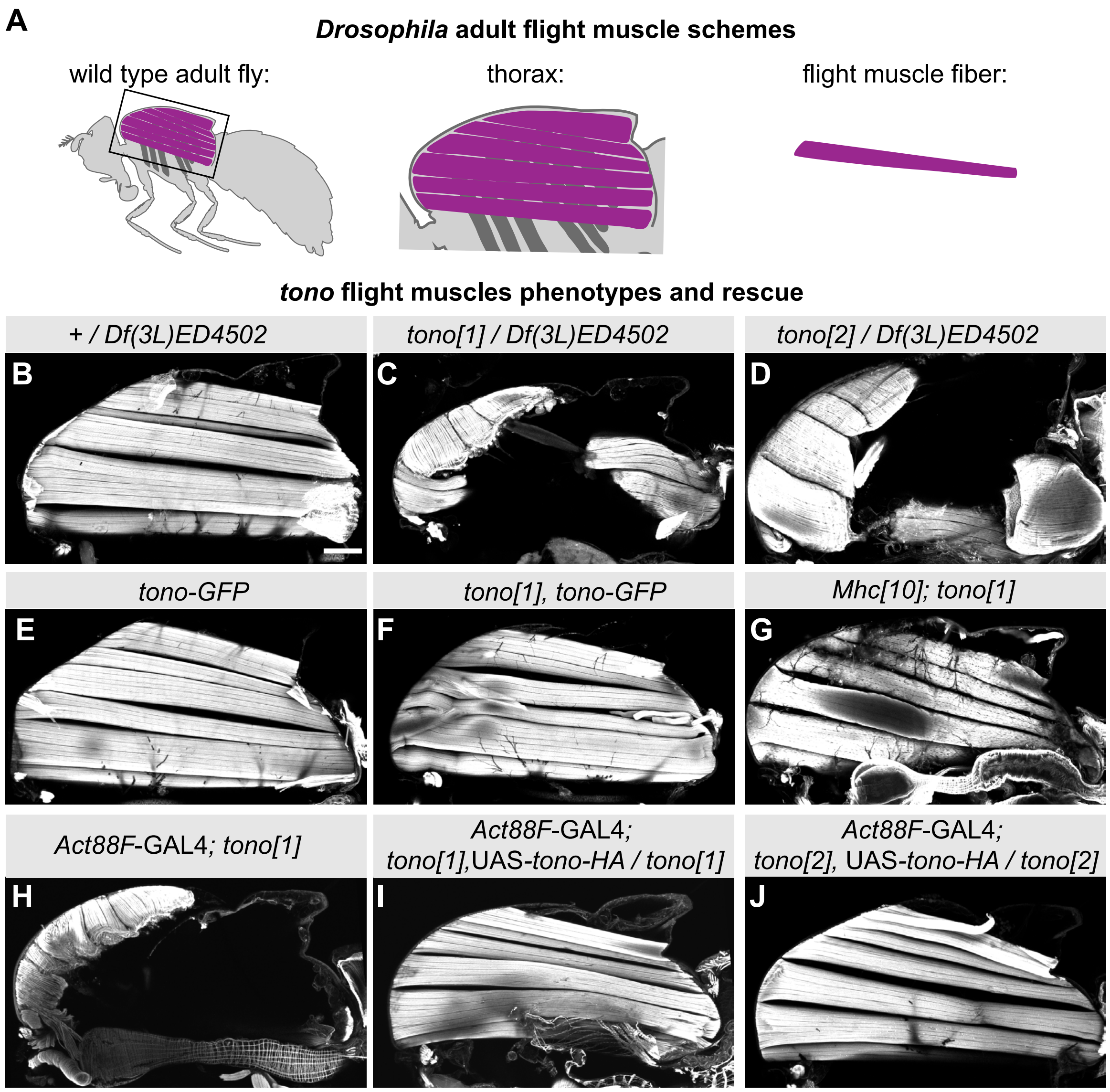


**Zhang et al. - Supplemental Figure 1**

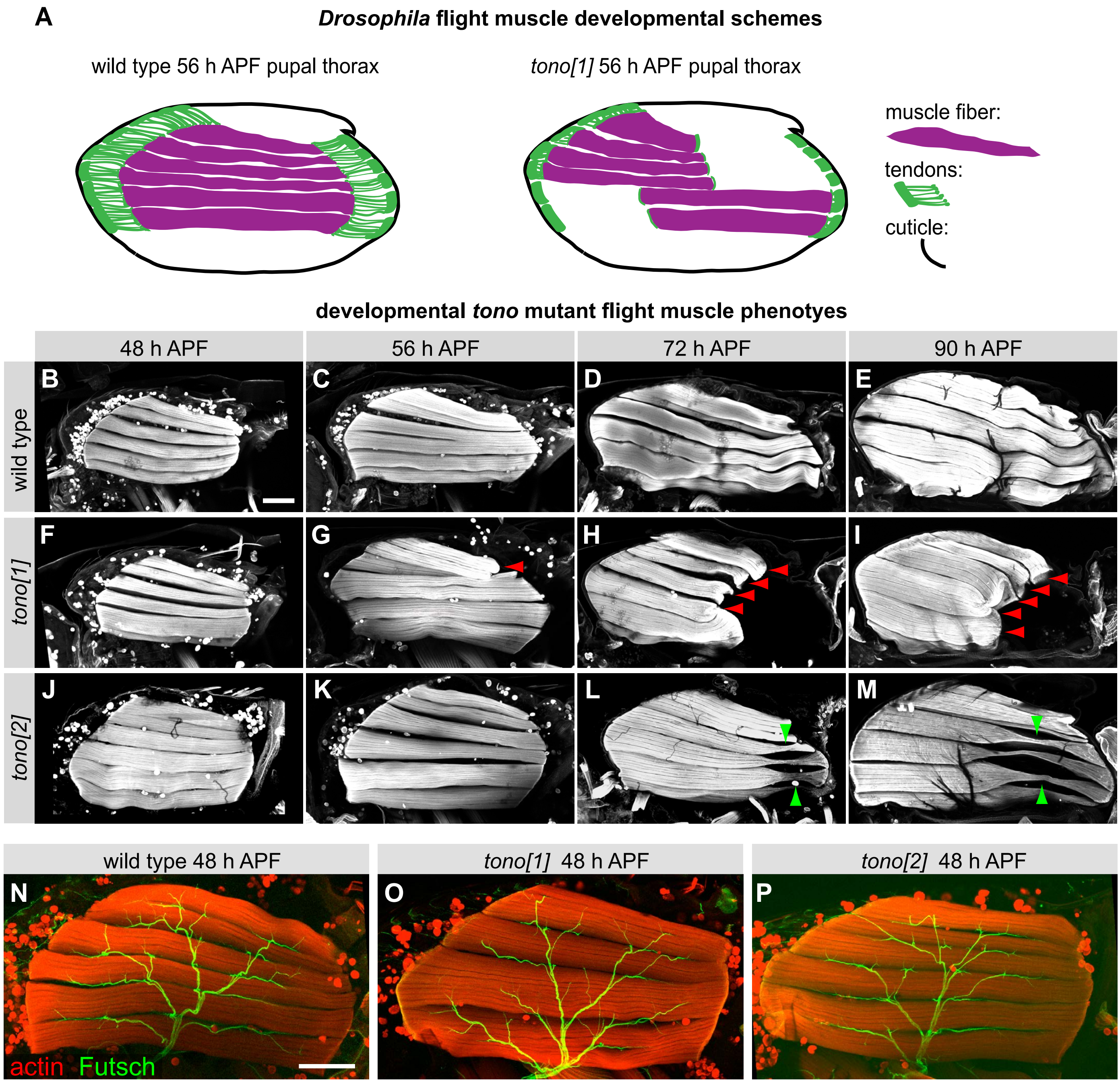


**Zhang et al. - Supplemental Figure 2**

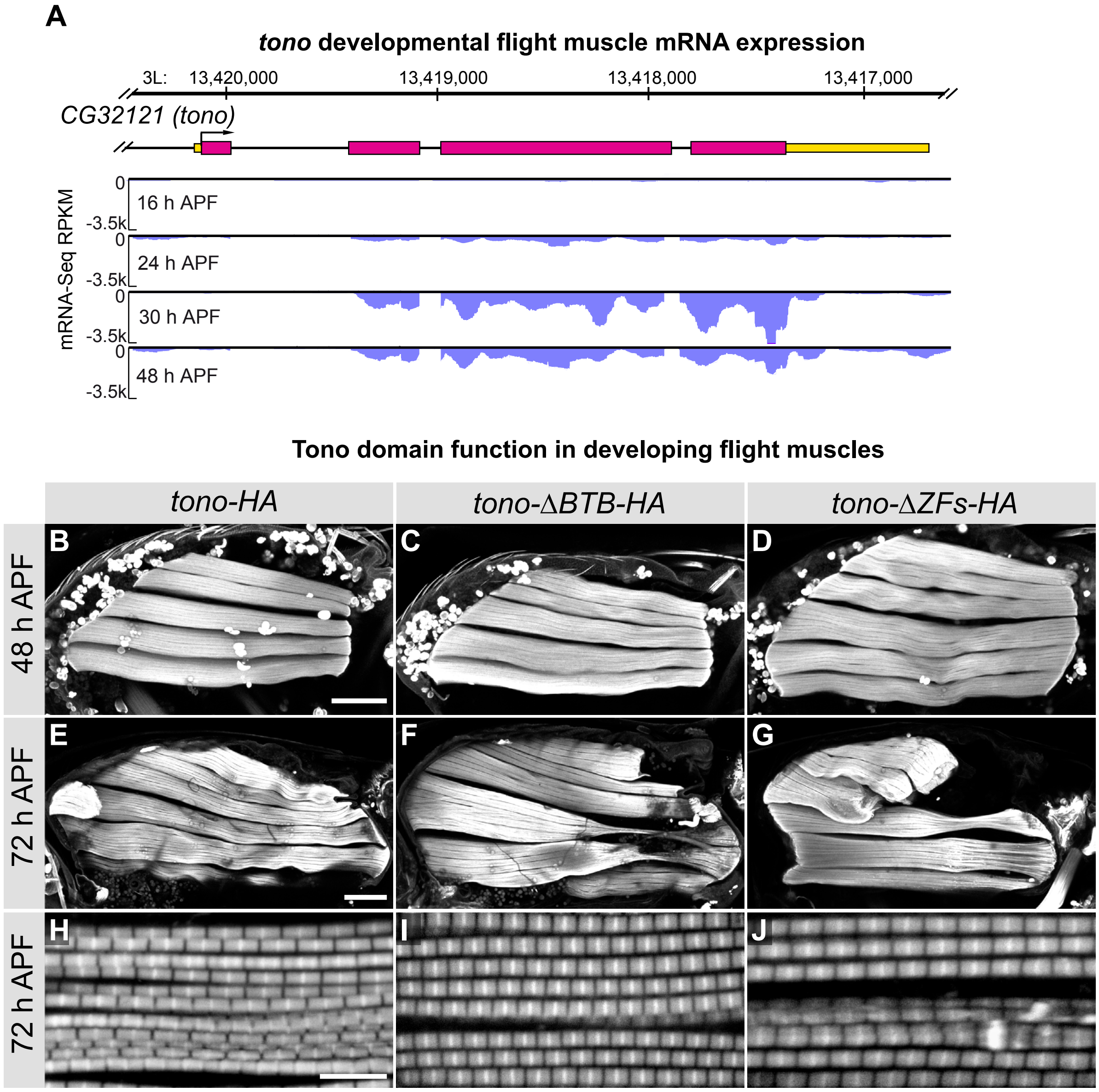


**Zhang et al. Supplemental Figure 3**

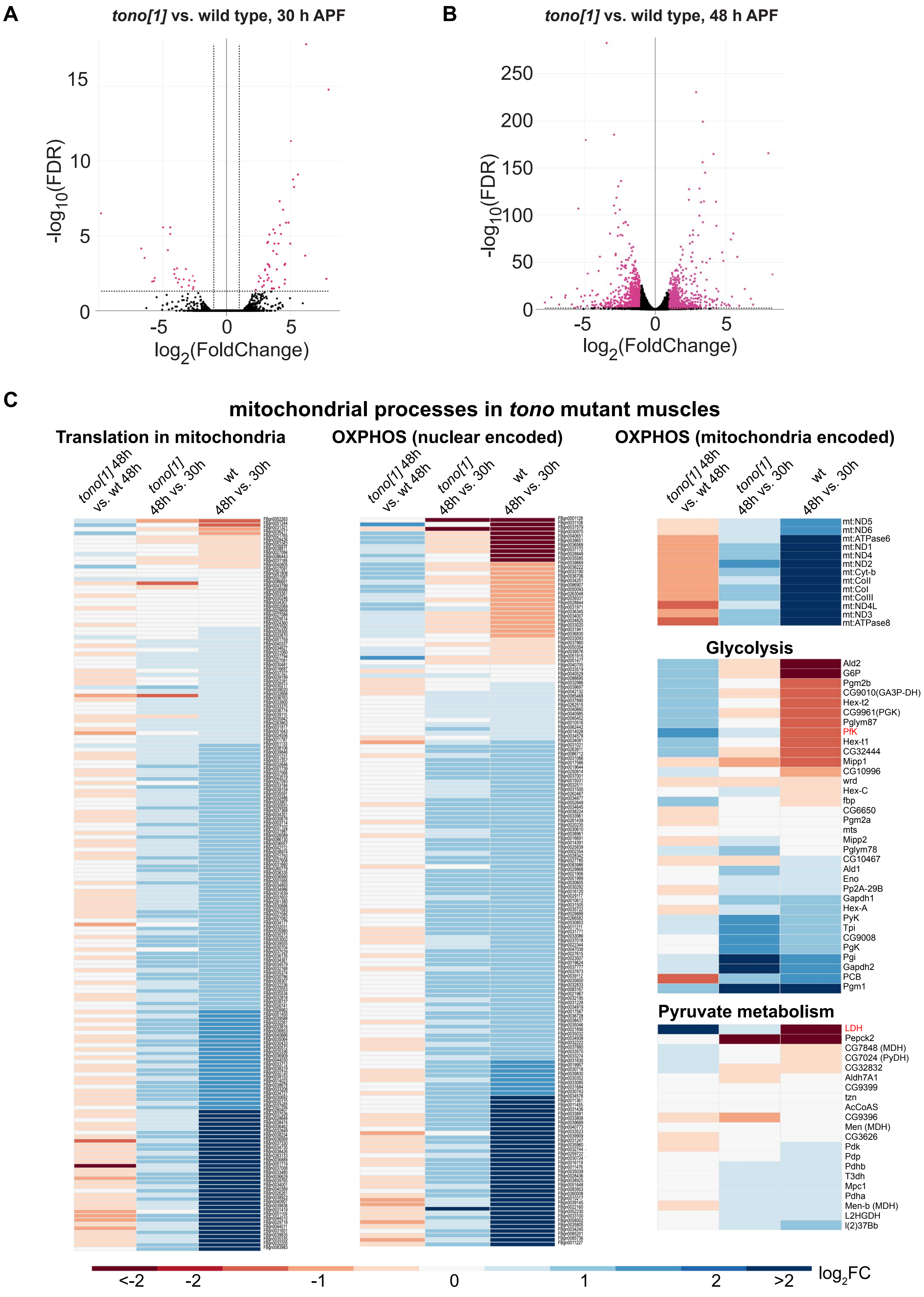


**Zhang et al. Supplemental Figure 4**

### 48 h to 30 h APF mRNA dynamics in wild type and *tono* flight muscles

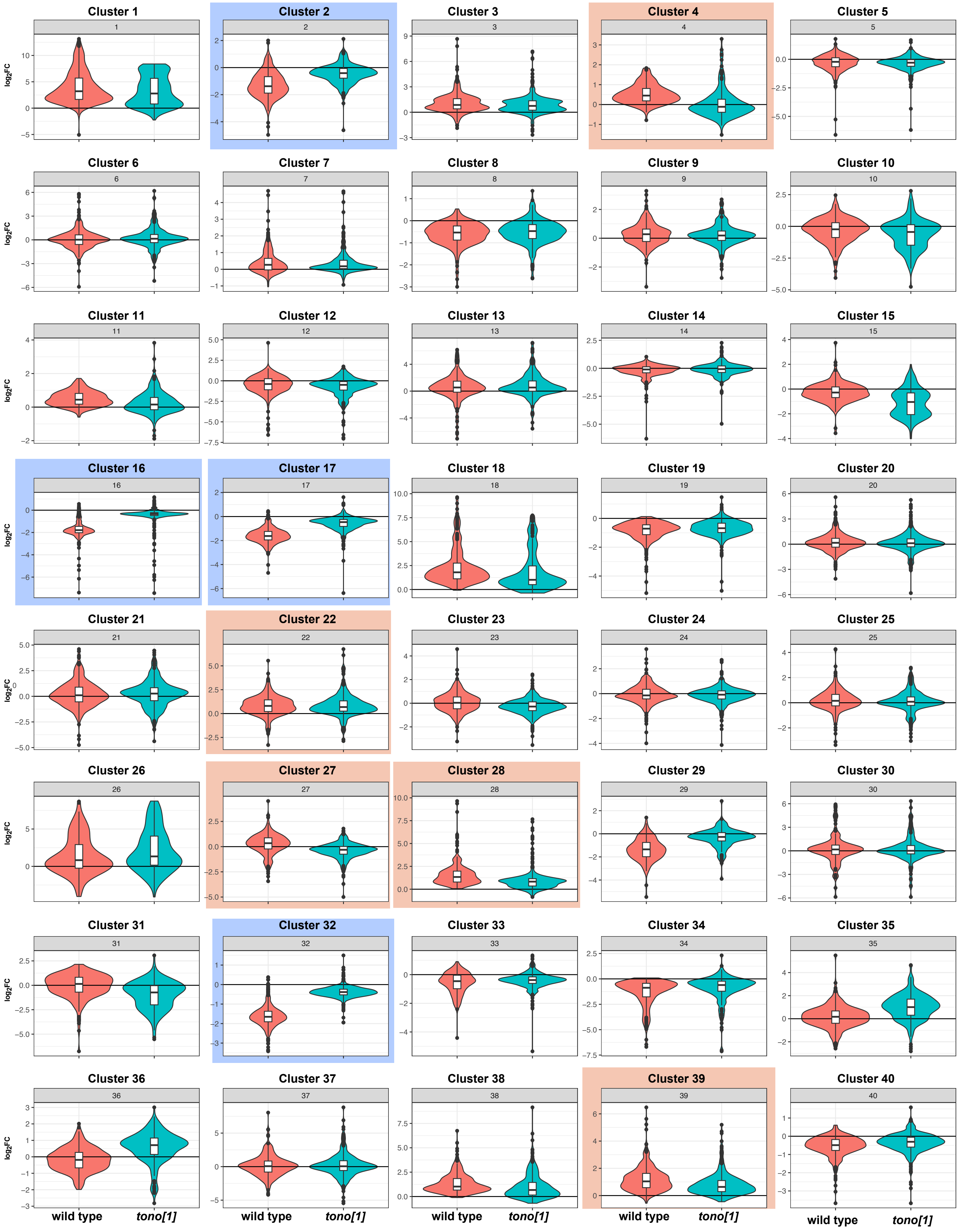


**Zhang et al. Supplemenal Figure 5**

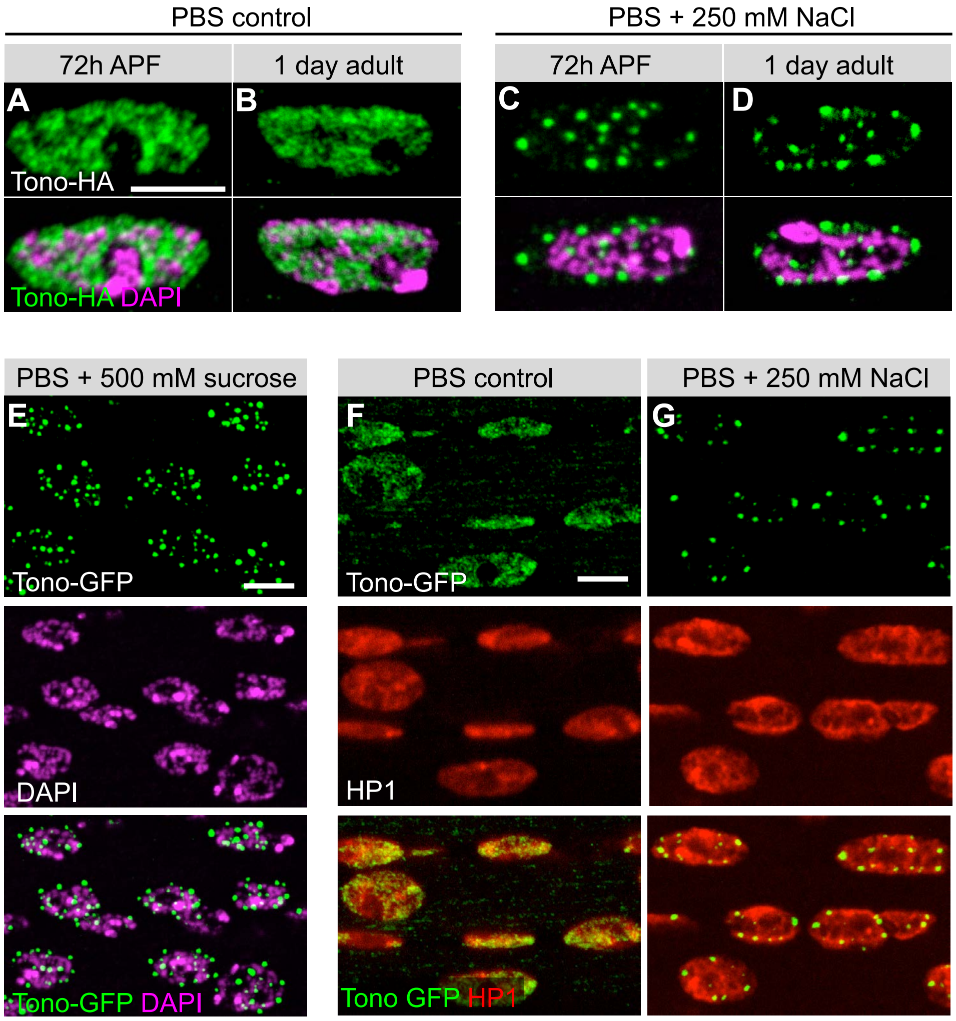


**Zhang et al. Supplemetal Figure 6**

#### Tono droplets are distinct from insulator bodies

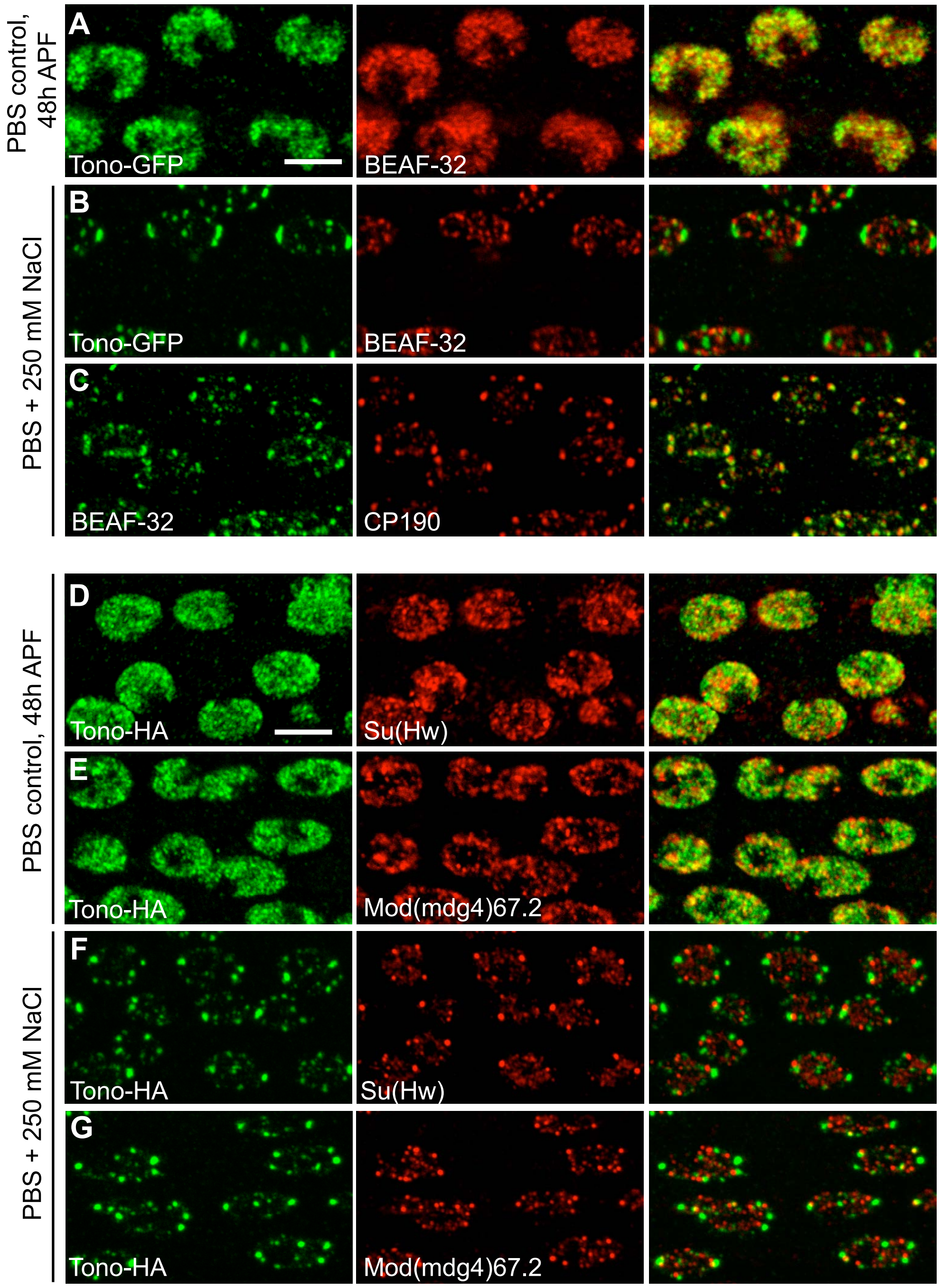


#### Insulator body protein mutant phenotypes

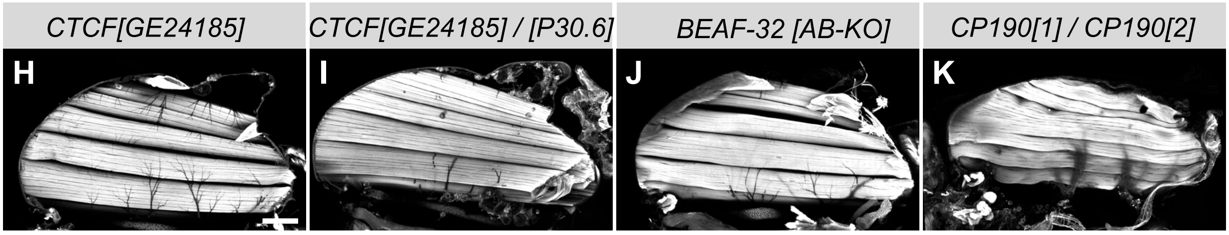


### Zhang et al. Supplemental Figure 7

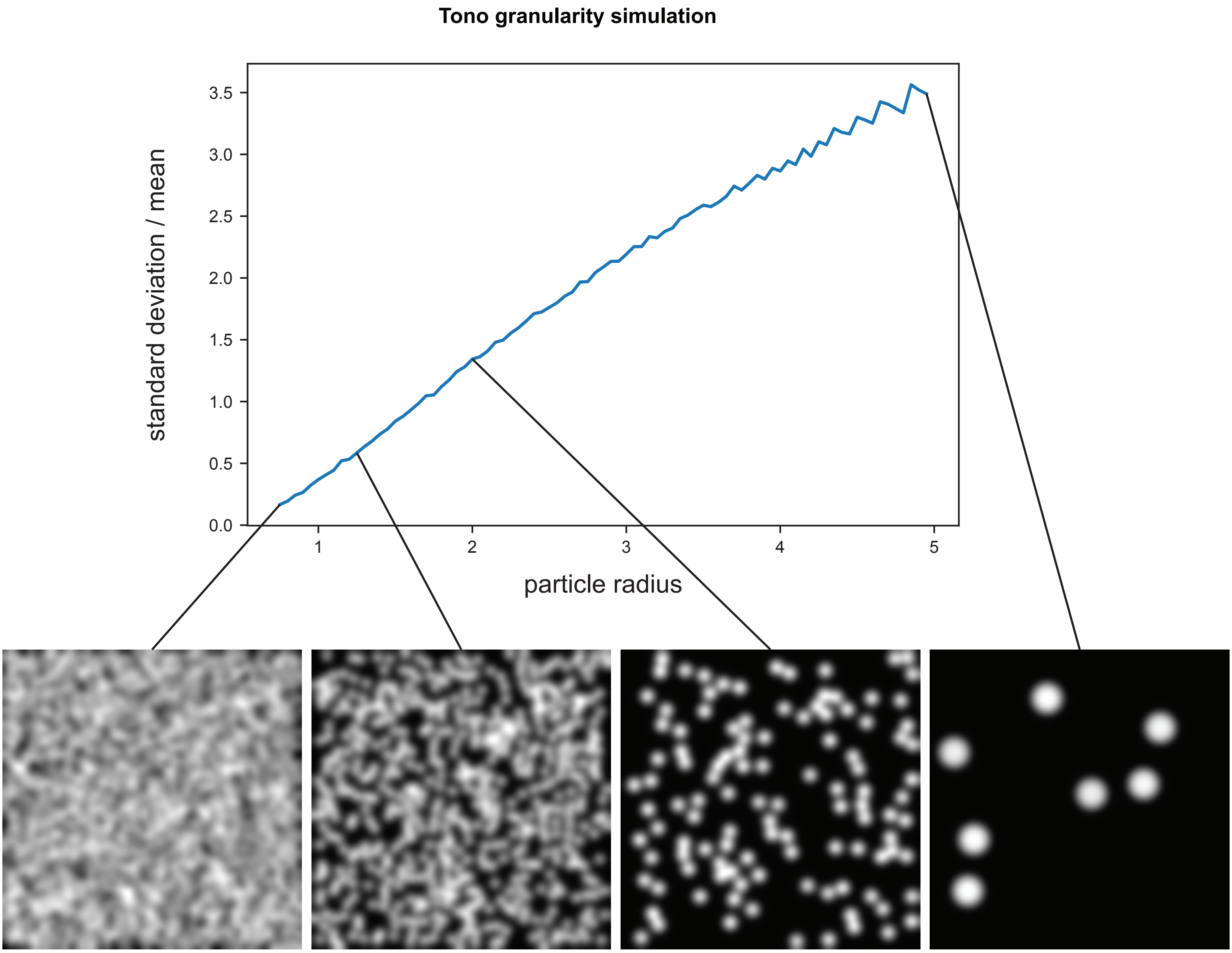


**Zhang et al. Supplemental Figure 8**
